# Modelling DNA replication fork stability and collapse using chromatin fiber analysis and the R-ODD-BLOBS program

**DOI:** 10.1101/2024.11.01.621594

**Authors:** Kerenza Cheng, Kazeera Aliar, Roozbeh Manshaei, Susan L Forsburg, Ali Mazalek, Sarah A Sabatinos

**Author notes:** correspondence, www.SabatinosLab.net.

## Abstract

We describe the anatomy of replication forks by comparing the lengths of synthesized BrdU-labelled DNA in wild type, *mrc1Δ* and *cds1Δ Schizoasaccharomyces pombe*. We correlated Rad51 and Cdc45 proteins relative to their positions on the fork, replicated tract, or unreplicated regions. We did this by using chromatin fiber images. These fibers track pixel intensity data, which is analyzed using our program: R-ODD-BLOBS. We compared the lengths of BrdU tracts and proteins, as well as the percentage of Rad51 and Cdc45 colocalization, and compared our results with literature findings. We measured average BrdU lengths consistent with current literature; *cds1Δ* was the longest at ∼2.9 kb (8.6 pixels, px), wild type was ∼ 2.5 kb (7.5 px), and *mrc1Δ* was the shortest at ∼1.7 kb (5.1 px). Intriguingly, Rad51 was found at 22% more replicated areas in *mrc1Δ* than in wild type. This suggests that homologous recombination repair may be more common at *mrc1Δ* forks. In this study, we summarize the usefulness of a computational modeling tool to assess large datasets of chromatin spread data. In turn, we find patterns of DNA replication length and protein components at replication forks, to describe the anatomy of a fork and how structures change with checkpoint loss.

GRAPHICAL ABSTRACT:
R-ODD-BLOBS uses chromatin fiber data to rigorously model replication fork structures.
DNA replication forks are multi-subunit structures that must pair and regulate DNA copying activity of the polymerases with unwinding activity of helicase. Chromatin fiber data retains proteins, and can be used to detect DNA synthesis (blue) and associated DNA replication fork proteins such as MCM4 helicase (MCM4) and replication protein A (RPA). In our work, we have used homologous recombination protein Rad51 and helicase factor Cdc45 to understand how DNA replication fork structures are destabilized during hydroxyurea treatment, and how they fail to recover because of Cdc45/helicase mis-localization.

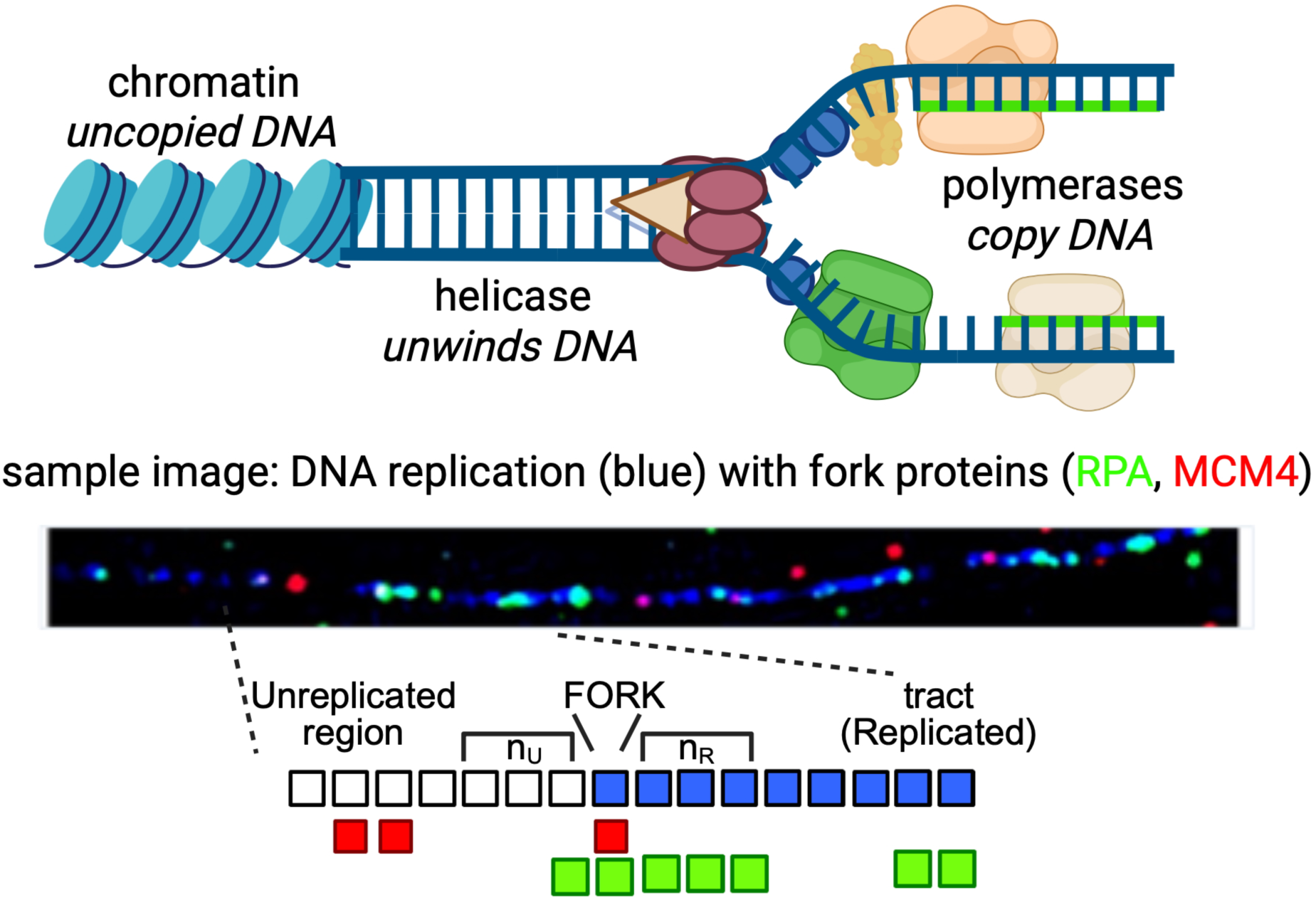

## 1.0 INTRODUCTION

High-fidelity DNA replication and active response to replication stress are critical to genome stability and cancer avoidance [1,2]. The replication fork complex consists of two linked activities that ensure DNA replication is efficient, speedy and faithful [3–6]. First, the Cdc45-MCM-GINS (CMG) helicase complex unwinds the DNA template ahead to promote high-speed copying of the genome [7,8]. Second, DNA polymerases on the leading (polymerase ε) and lagging (polymerases Δ and ɑ) strands create two copies of DNA to be split between daughter cells at division [9–11]. The Claspin/Mrc1 protein is a part of the Fork Protection Complex that directly connects polymerases to the helicase complex (*e.g.* [12,13]). Claspin/Mrc1 also influences replication processivity and independently activates Cds1 (mammalian CHEK1) for the replication checkpoint [13–17].

The replication checkpoint responds to various naturally occurring and exogenous obstacles that cause replication fork slowing and/or stalling [18]. A drug that causes DNA replication arrest is hydroxyurea (HU). HU depletes deoxynucleotide triphosphate (dNTP) pools and causes forks to slow and arrest [19–21]. In the fission yeast *Schizosaccharomyces pombe*, the checkpoint protein Cds1 is phosphorylated and activated by Rad3 kinase (ATR homologue). Activated Cds1 promotes stable replication arrest in the presence of HU as dNTP levels decrease [22]. Cds1-mediated arrest prevents late-origin firing, rescuing the remainder of the DNA replication program [21]. In a *cds1Δ* mutant, cells continue to synthesize DNA in HU and acquire DNA damage [21,23]. This leads to *S. pombe* cell elongation and a lethal G2 arrest.

Replication fork stalling and collapse generates DNA damage. Double-strand DNA breaks (DSB) are a particularly lethal form of DNA damage, that is known to occur after DNA replication instability. DSBs may be repaired by homologous recombination (HR) [24,25], a pathway that is initiated by the Mre11-Rad50-Nbs1 (MRN) complex. MRN and co-factor CtIP recognize a DSB end and resect the break, allowing Rad51 recruitment to the resulting single-stranded DNA (ssDNA) [25]. Rad51 is activated by mediators, including Rad52, allowing Rad51 to search for homologous regions that serve as a template for repair [25,26].

Replisome dynamics during HU arrest have been modeled in molecular studies, where immune-precipitated complexes are assessed. The results from these whole-cell population lysates is averaged across a population using sequencing, or isolation of proteins on nascent DNA (iPOND) [27–29]. Consequently, our model of replisome arrest in HU considers bulk protein movement and association with specific sequences. However, the average response across a population may obscure some events through “ensemble averaging” [30,31]. Thus, the behaviour of sub-populations with alternate phenotypes is lost [32], making it impossible to understand the diversity and range of phenotypes present within a cell. Understanding the diversity of replisome effects would allow us to model replisome stability mechanisms; a detailed understanding of replication fork anatomy and variation would help us to understand how instability and mutation occur. Other experiments using electron microscopy have also imaged DNA replication forks and detected increased ssDNA at individual forks during arrest and replicon components show association and structure, but in the absence of genome context and replisome protein components [11,33].

Our microscopy and molecular labeling approach on chromatin fibers allows a deeper exploration of replication dynamics within the genome. The chromatin fiber analysis is an inexpensive and high-content method that retains proteins, enabling the direct detection of replication complexes on DNA with fluorescent tagging. A key barrier to using the data is in the extensive imaging data that is generated; analysis and modeling is a challenge. We developed a program called One Dimensional Data – Boolean Logic Operations Binning System (ODD-BLOBS) to systematically correlate DNA synthesis with protein location using chromatin fiber data [34]. ODD-BLOBS measures tract lengths to determine replication processivity or instability. In this work we describe a new implementation of ODD-BLOBS in the programming language R (now R-ODD-BLOBS). R-ODD-BLOBS is robust and user-friendly, allowing quick processing of linear data to identify replication tracts, isolate replication fork zones, smooth gaps in signal, and highlight co-localized replisome components [34].

We have compared fibers from wild type (wt), *mrc1Δ* and *cds1Δ S.pombe* cultures labeled with bromodeoxyuridine (BrdU). BrdU detects newly synthesized DNA following HU arrest. Cdc45 and Rad51 proteins are described relative to synthesis to describe how replication checkpoint protects the genome in response to HU. The mean lengths we observed using R-ODD-BLOBS accurately reflect the condensed and filamentous nature of the DNA and proteins. Cdc45 and Rad51 colocalization determined by R-ODD-BLOBS potentially can be used to detect and predict HR. The *mrc1Δ* cells in particular show different Rad51 patterns. By standardizing chromatin fiber data analysis methods, we correlate protein position relative to synthesis in the visual anatomy of the replication fork. This new R-ODD-BLOBS method of analysis establishes a single-molecule approach to study replication fork dynamics.

## 2.0 MATERIALS & METHODS

We used wildtype, *mrc1Δ,* and *cds1Δ* yeast cell strains that express the transgene of *hsv-tk* and *hENT1* for the experiments (as previously described in [22,23]). Yeast cultures were grown in pombe minimal glutamate media (PMG with 225 mg/mL of adenine and uracil) to mid-log phase (i.e., OD600 of 0.4 to 0.8, approximately 1 × 106 cells/mL). Hydroxyurea (HU, BioShop, Canada) was added to 12 mM for 4h at 30°C with shaking. HU arrests cells in S-phase. Cells were collected onto whatman vacuum filters, and washed twice with pre-warmed medium. Cells were resuspended in pre-warmed medium, and 50 µg/mL BrdU was added for 30 minutes.

Chromatin fibers were prepared as described in [34]. Briefly, cells were harvested, centrifuged, and washed with water. Pelleted cells were resuspended in zymolyase mixture (1 *M* sorbitol, 60 m*M* EDTA, 100 m*M* sodium citrate, 0.5 mg/mL zymolyase 20T, 1.0 mg/mL lysing enzymes, 100 mM 2-mercaptoethanol, pH 6.9) for 15 to 30 min. Spheroplasting was checked by microscopy. Spheroplasts were centrifuged gently, and then resuspended in 500 µL of phosphate buffered saline.

Spheroplasts were adhered onto poly-lysine coated coverslips (number 1.5 thickness). Lines were incubated with pre-warmed lysing buffer for 15 min (50 m*M* Tris-HCl pH 7.4, 25 m*M* EDTA, 500 m*M* sodium chloride, 0.1% Nonidet P-40 (Sigma), 0.5% (w/v) sodium dodecyl sulphate (SDS, BioShop), 3 m*M* 2-mercaptoethanol; 70°C). Fibers were stretched at a 10° to 25° angle, and air dried. Fibers were fixed in 4% paraformaldehyde in phosphate buffered saline (pH 7.4) for 10 minutes, before being rinsed in PBS and then baked for 5 minutes at 60°C. Samples were stored at -20°C until detection.

### 2.1 Fiber labeling methods

To detect BrdU, fibers were first wet in PBS and then denatured in 2 N HCl for 15 minutes. Samples were then neutralized in 0.1 *M* sodium tetraborate (Na_2_B_4_O_7_; BioShop Canada) solution (pH 8.5) for 10 minutes, and then washed in PBS. Slides were blocked using 10% fetal calf serum, 10% bovine serum albumin, 0.05% Tween 20 detergent in PBS and filter sterilized in a dark humid chamber. BrdU was detected with rat BU1/75 (ICR1, Abcam), diluted to 1:100 in blocking buffer. Cdc45-myc was detected with anti-myc antibody (9E10 clone, Covance), FLAG -polymerase alpha with anti-FLAG M2 (Sigma), and Rad51 rabbit anti-Rad51 (Forsburg lab). Primary antibodies were incubated overnight at 4°C in the dark. After washing in PBS three times, secondary antibodies were applied and incubated at 30°C for 2h. Secondary antibodies against each primary antibody were diluted in blocking buffer at 1:500, including chicken anti-rat AlexaFluor 488 (Invitrogen), goat anti-rabbit AlexaFluor 546 (Invitrogen) and donkey anti-mouse AlexaFluor647 (Invitrogen). Samples were washed in PBS, and mounted onto glass slides with SlowFade Gold antifade mount containing 1 µg/mL 4’,6-diamidino-2-phenylindole dihydrochloride (DAPI) (Invitrogen).

Samples were imaged using a DeltaVision microscope with softWoRx v4.1 (GE, Issaquah, WA) using a 60x (NA1.4 PlanApo) lens, solid-state illuminator and 12-bit CCD camera. Images were acquired in five 0.2µm z-sections, then deconvolved and Maximum Intensity Projected (softWoRx, default settings). Image stacks were deconvolved and projected. Fibers were manually traced by drawing line segments with the “Arbitrary Line Profile” tool (SoftWorx, Applied Precision Instruments). Fluorescent intensity values from each channel relative to pixel location were acquired along these lines. Data was processed into tab-delimited files for analysis with ODD-BLOBS or R-ODD-BLOBS. A minimum of 3 fields per strain was analyzed, and data was compared among experimental replicates.

We ran the intensity data in R-ODD-BLOBS to obtain the initial scatter plot data and determine the preliminary baseline threshold values. We re-ran R-ODD-BLOBS by fixing 2 channels with baseline threshold values, and iterating the dependent variable from thresholds of 100-1000, increasing by increments of 100 (all fluorescent intensity measurements are in arbitrary units, AU). We found that optimal thresholding values for our dataset were DNA at 100, and BrdU, Rad51 and Cdc45 is 200. After thresholding was confirmed, we used a smoothing effects analysis to test smoothing values from 0-6, with optimal threshold values found above. The best smoothing value for our data was 3 for DNA and BrdU, and 4 for Rad51 and Cdc45. After the threshold and smoothing were determined, R-ODD-BLOBS was run using the optimized values to determine protein colocalization within each region (*i.e*. unreplicated, fork, replicated) and to calculate length of BrdU or Cdc45/Rad51 tracts.

## 3.0 RESULTS AND DISCUSSION

We first assessed chromatin fiber characteristics from microscopy images. We tested the length of chromatin stretching in wild type fibers. A fluorescence *in-situ* hybridization (FISH) cosmid probe was hybridized to fibers. The 28.6kb cosmid probe c1095 is on chromosome 1 (0.726-0.755 Mb) within 10 kb of an early and strong origin, ORI 12 [35–37]. We found a mean of 9.2 µm per probe signal, indicating approximately 3.1kb/µm of DNA extension (standard deviation 2.12 kb/µm) (Figure 1A, 1B). With a pixel size of 0.1092 µm/pixel in our microscopy system, this converts to approximately 331 bp/px in our images. With this imaging (px) to DNA (bp) conversion value in mind, we continued to analyze replication tracts and associated factors using pixels as the smallest but easily convertible unit of measurement.

**Figure 1.**
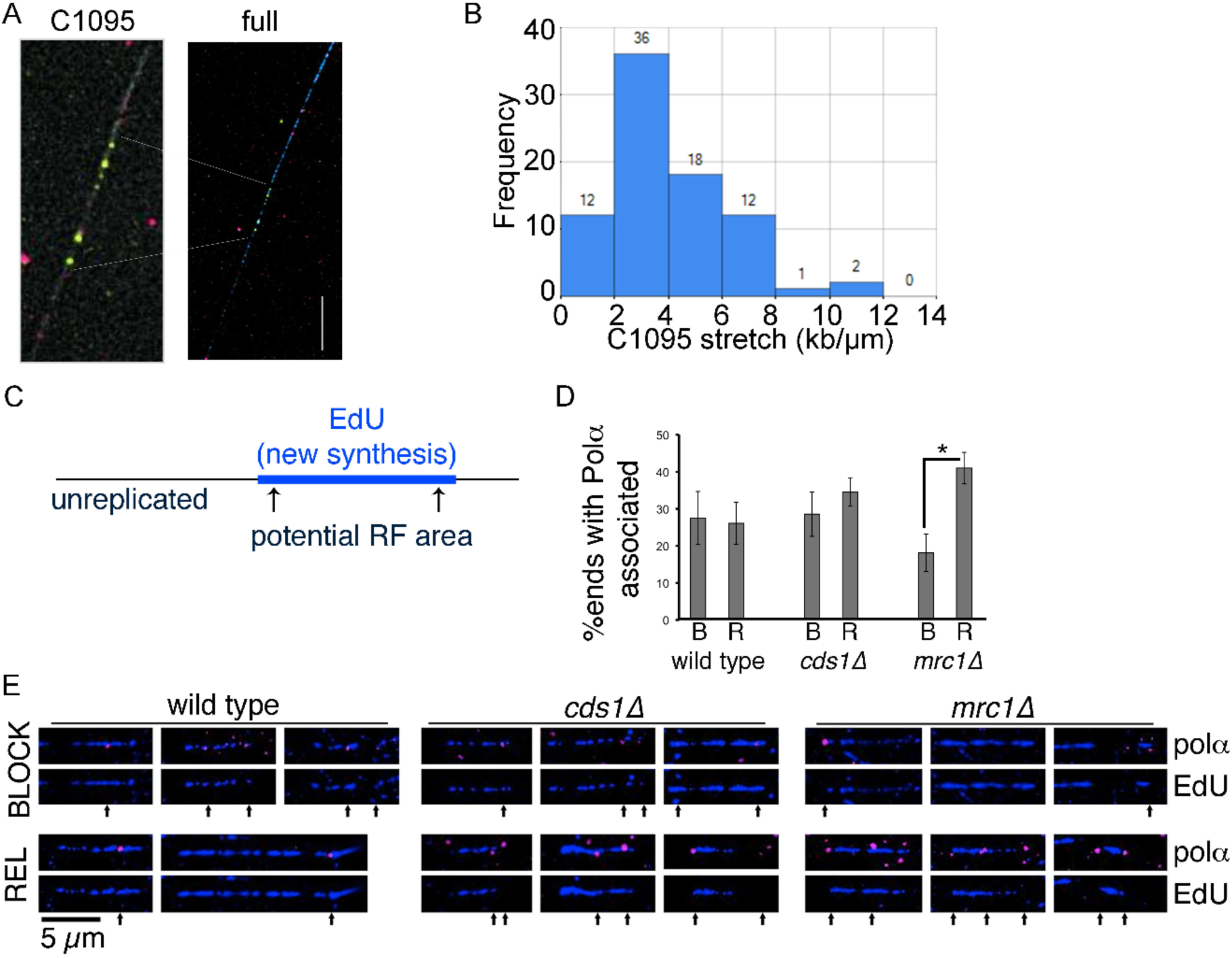
Data analysis of wt, *cds1Δ,* and *mrc1Δ* on polα using ODD-BLOBS. Data analysis using ODD-BLOBS code and replication fork detection principles. (A) A fluorescent microscopy image of a chromatin spread fiber with a fluorescence in-situ hybridization (FISH) cosmid probe to detect c1095 area of the genome. BrdU label is blue, MCM2 helicase component is red, and c1095 FISH probe is green. A c1095 area is highlighted, showing the signals used to calculate stretching in B. (B) Histogram of C1095 distances, converted to kb/µm. Data from n=81 c1095 tracts, over 3+ images and samples, was measured in pixels from end to end. c1095 tracts were converted to kb/µm using the known c1095 distance of 28.6kb. Fiber stretch centers between 2-4 kb/µm, and the mean value for fiber stretch was calculated at 3.1 kb/µm. (C) Scheme for analyzing replicated tracts on spread DNA fibers. Each replicated tract has an internal (replicated) zone and external unreplicated areas. We defined the ends of each synthesized tract as potential replication fork areas. Bi-directional replicons would have 2 forks. (D) Newly synthesized tracts were detected using EdU labeling and then Click chemistry. DNA primase (polα-FLAG) was detected using with anti-FLAG and fluorescent secondary antibody detection, relative to EdU incorporation and tract-ends (potential replication fork areas). EdU ends with Polα are presented as the percent of total EdU tract ends examined during HU block (labelled B) or following release from drug (labelled R). Proportion shown with 95% confidence interval from 2+ replicates. (E) Microscopy images of Polα (pink) at EdU tract-ends (tag in blue) during HU (BLOCK) and release (REL) as quantified in (D). EdU tracts only are shown below merged image with Polα in potential replication fork areas indicated by arrows under the tracts.

### 3.1 Polymerase alpha (α) associated with replicated tracts in checkpoint mutants

We have seen that *cds1*Δ and *mrc1*Δ mutants continue DNA synthesis during and after HU arrest [23]. Now, we asked whether replication fork components remain associated with the fork in the absence of Cds1 or Mrc1. Chromatin DNA fibers retain proteins in a chromatin setting [38], even though some protein is lost during preparation. To minimize the impact of non-fiber protein detection, we tested samples without primary antibody and by switching fluor (excitation/emission) combinations (data not shown). We used a FLAG antibody to detect FLAG-polymerase α on fibers. The fibers were labeled with the thymidine analogue EdU to mark new DNA synthesis (Figure 1C). We defined the tips of EdU tracts as potential replication fork locations, as described for ODD-BLOBS [34], and detected FLAG-Polα either on or near EdU-tracts.

Because HU arrests cells in S-phase, we compared polymerase alpha and EdU tracts after 4h in HU, or at 30 minutes after HU removal. We found similar polymerase alpha associated with EdU tracts during block or release in both wild type and *cds1Δ* (Figure 1D, 1E; FLAG-Polα are pink foci on blue EdU tracts). However, the *mrc1*Δ tracts had less tract-associated polymerase alpha during the HU block. Once HU was removed, we saw that *mrc1*Δ tracts regained Polα like wild type and *cds1*Δ. Our analysis confirmed that proteins were enriched near replicated tract ends. Confirming that proteins were located on chromatin fibers and near replicated tracts formed a basis for next steps of computational analysis using R-ODD-BLOBS.

### 3.2 Thresholding analysis using R-ODD-BLOBS

The previous ODD-BLOBS software to analyze chromatin fibers used visual basic for applications (VBA) script [34]. While ODD-BLOBS ran in Excel and Libre Office platforms, the interface was difficult to use and processing was time-intensive. We adapted ODD-BLOBS into R-script to make R-ODD-BLOBS.

R-ODD-BLOBS first creates an initial scatter plot of the initial intensity values (Supplementary Figure *S1)*. We used scatter plots to determine a preliminary baseline threshold based on the distribution of signal intensity for each channel. Outlying data points are filtered using these low-level methods, to prevent outliers from skewing later thresholding results. A preliminary baseline threshold was determined as a placeholder for iterative testing of each channel: 150 for BrdU, 120 for Rad51, and 130 for Cdc45.

We next tested threshold intensity values from 100 to 1000 in each channel. A general trend found in all channels and conditions of thresholding is that as the threshold increases, the length of the tracts get shorter. For example, as BrdU threshold increases, fewer pixels are above threshold values, and the length of tracts detected is decreased because of gaps in signal or sub-threshold values (Figure 2A). We found that there is a point of saturation where the effect of threshold increase is minimal. The tract length of BrdU in wildtype shows a slight decrease as the threshold gets larger, but plateaus after a threshold of 300 (Figure 2A). At the lowest threshold of 100, some tracts are over 390px in *cds1Δ* and *mrc1Δ* (Figure 2B, 2C). However, we still observed a saturation point in threshold to length ratios, above which the change of threshold value is minimal. This observation was consistent between replicate sample populations (Supplementary Figure S2).

**Figure 2:**
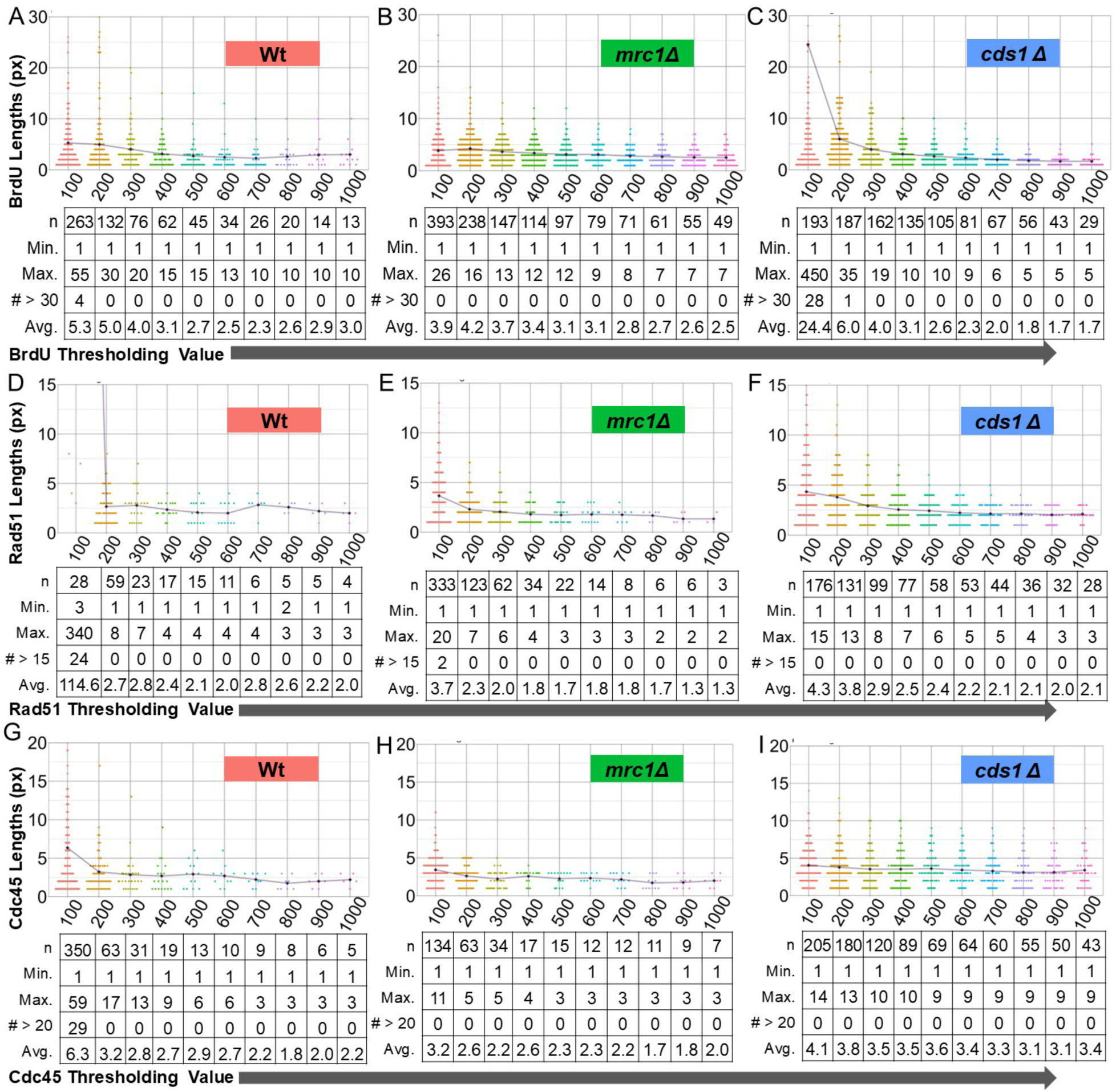
Increased fluorescent threshold values decrease BrdU, Rad51, and Cdc45 lengths. The intensities of the line-traced chromatin fiber data was processed using R-ODD-BLOBS using different thresholds for each channel. Thresholds from 100 to 1000 were used to model the effect of higher threshold on tract length of DNA synthesis (BrdU) or replication associated protein (Rad51, Cdc45). Each channel was iterated separately. A baseline “placeholder” threshold was used for each channel not tested. Baseline thresholds were calculated using a scatter plot of intensities, shown in Supplementary Figure S1 as: 100 for DNA, 150 for BrdU, 120 for Rad51, and 130 for Cdc45. Note that chart scale is set from 0 to 30 for tract length; outliers are described with maximum values in the chart below each threshold bin. (A-C) Violin plots of the tract lengths in BrdU in wt, *cds1Δ*, and *mrc1Δ* at thresholds from 100-1000. (D-F) Violin plots of the protein lengths in Rad51 in wt, *cds1Δ*, and *mrc1Δ* at thresholds from 100-1000. (G-I) Violin plots of the protein lengths in Rad51 in wt, *cds1Δ*, and *mrc1Δ* at thresholds from 100-1000. The orange triangle on the plots represents cut-off data at that threshold.

While increased threshold values tended to decrease tract length, we found that Rad51 tracts were small and least impacted by threshold (Figure 2D, 2E, 2F; Supplementary Figure S3). Average Rad51 tracts of any genotype did not change above a 300 fluorescent intensity-threshold setting. Curiously, we found that Cdc45 protein length distributions (Figure 2G, 2H, 2I) had bi-modal groupings in replicate data sets below threshold of 500 (Supplmentary Figure S4). Generally, the average tract length of a protein or BrdU decreases as thresholds increase. However, bimodal populations uniquely show a slight increase in mid-range thresholds that increase with higher threshold. For example, Cdc45 in wild type (Figure 2G) has a bimodal population at a threshold of 500. Cdc45 in *mrc1Δ* (Figure 2I) shows a longer population of tracts at threshold values of 400-600.

### 3.3 Smoothing extends tracts and minimizes signal gaps

Small pixel gaps between signals can be smoothed with the R-ODD-BLOBS smoothing function. A 3 pixel or smaller gap in signal cannot be confidently resolved because of the Abbé Limit using our system. R-ODD-BLOBS can model data with small gaps joined, which may correct missing signal in datasets. Importantly, we can compare R-ODD-BLOBS results with and without smoothing to model the impact on tract lengths (Figure 3) and protein associations (Figure 6). We used threshold values optimized in Figure 2 for BrdU, Rad51, and Cdc45. We then iteratively smoothed pixel gaps from 0 to 6 px (between above-threshold areas) on each channel (BrdU, Rad51, and Cdc45). This was done using the “Smooth-It” parameter in R-ODD-BLOBS, similar to the method described in [34].

**Figure 3:**
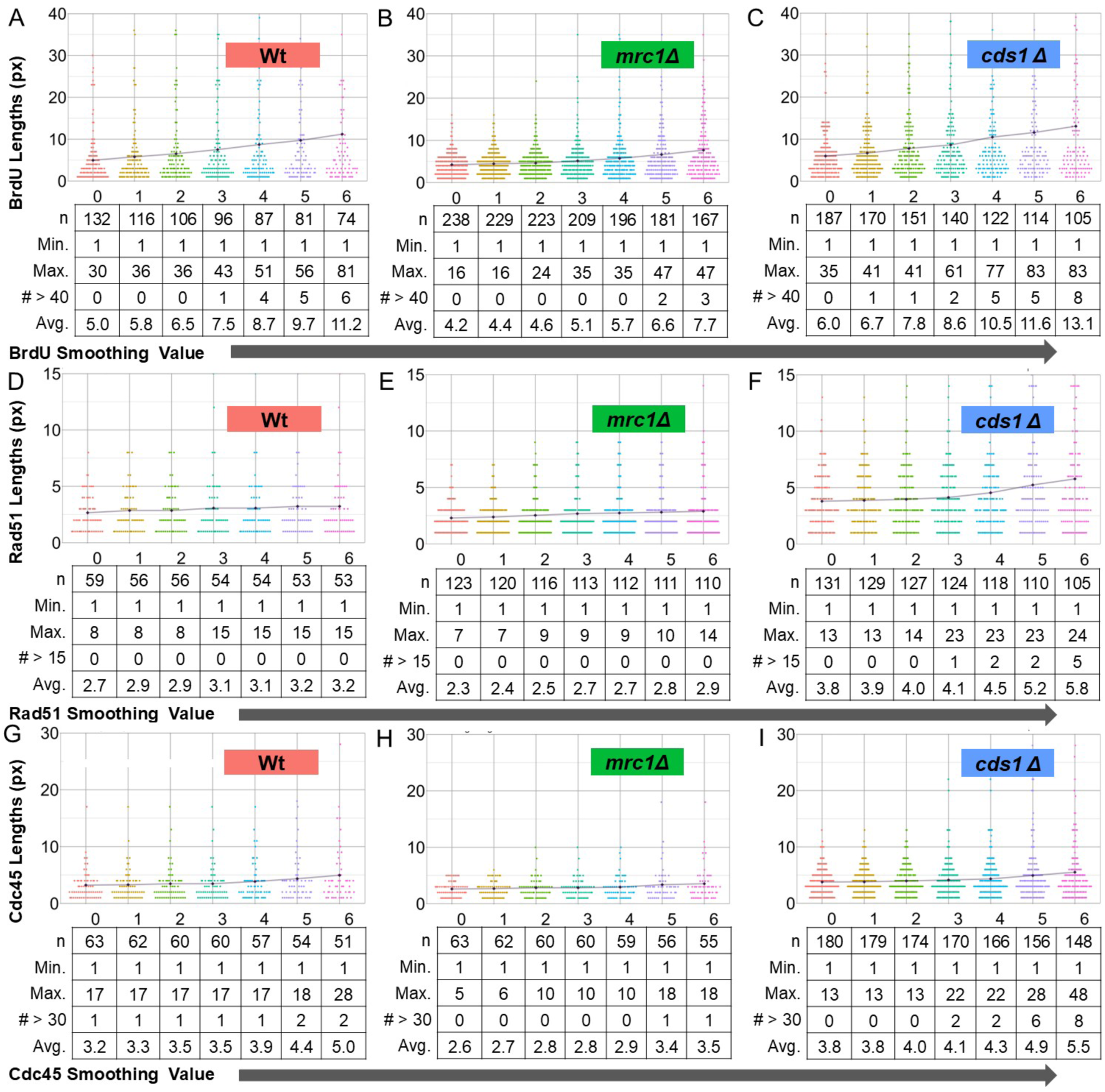
Larger smoothing values contribute to longer tract lengths for all of BrdU, Rad51, and Cdc45. The intensity of the line-traced chromatin fiber data was processed using R-ODD-BLOBS using previously determined threshold values: 100 for DNA; 200 for BrdU, Rad51, and Cdc45. Smoothing links tracts together if the number of pixes is equal to or less than the “smoothing value” used. (A-C) Violin plots of the tract lengths in BrdU in wt, *cds1Δ*, and *mrc1Δ* at smoothing values 0 px to 6 px. (D-F) Violin plots of the tract lengths in Rad51 in wt, *cds1Δ*, and *mrc1Δ* at smoothing values 0 px to 6 px. (G-I) Violin plots of the tract lengths in Cdc45 in wt, *cds1Δ*, and mrc1Δ at smoothing values 0 px to 6 px.

We found that smoothing the gaps in signals generally increased tract lengths, regardless of identity (BrdU, Cdc45, Rad51) or fluor emission spectrum. Smoothing had a large effect on wild type and *cds1Δ* BrdU signals (Figure 3A, 3C), where the impact of smoothing 6 pixel gaps doubled average tract lengths. In *mrc1Δ* tracts the impact of smoothing BrdU signal was less dramatic (Figure 3B). Similarly, Cdc45 signal length was unchanged by smoothing gaps in signal (Figure 3G, 3H, 3I), consistent with Cdc45 signal as a single protein source that is not predicted to spread.

We hypothesized that Rad51 might spread around unstable forks to allow Rad51 filament formation in homologous recombination, fork regression, or to protect excess ssDNA [39–42]. Intriguingly, we found that Rad51 signal smoothing led to longer tracts in *cds1Δ,* but not in wild type or *mrc1Δ* (Figure 3D, 3E, 3F). Longer Rad51 tracts in *cds1Δ* are consistent with reports that Rad51 is regulated by Cds1 during HU arrest and restart [40]. Our data show that smoothing is appropriate to cover gaps in protein signal that would otherwise create more protein detection sites. Further, smoothed Rad51 tracts may be used to model ssDNA dynamics by bridging signals that are adjacent but not coincident. Smoothing tract lengths also limits multimodal distribution patterns.

### 3.4 Aggregate tract patterns show that synthesis is coincident with Rad51 in cds1Δ

We used R-ODD-BLOBS to calculate average tract lengths in each channel and compare between genotypes. This ensemble averaging estimates the strongest trend within a population and is analogous to iPOND or ChIP-sequencing results. BrdU lengths were statistically different between genotypes. The *cds1Δ* cells had the longest tracts at 8.6 px (approximately 2.9 kb). Wild type average tracts were 7.5 px (∼2.5 kb), while *mrc1Δ* had the shortest tracts at 5.1 px (∼1.7 kb) (Figure 4A). Based on thresholding and smoothing patterns, we hypothesized that Rad51 tracts would be longest in *cds1Δ* cells. We found that *cds1Δ* tracts of Rad51 are longer than either wild type or *mrc1Δ,* and extend an average of ∼1.5 kb (4.5 px, Figure 4B). We also found that Cdc45 signal was longer in *cds1Δ* (Figure 4C) with an average of 4.3 px (∼1.5 kb). This observation wt had a similarly long length at 7.5 px and *mrc1Δ* had the shortest lengths at 5.1 px (Fig 4B). In Fig 4C, the lengths of Rad51 in each were statistically different, Rad51 in *cds1Δ* had the largest difference and was also the longest at 4.5 px. Rad51 in *mrc1Δ* was the shortest and was nearly half the lengths of *cds1Δ* at 2.7 px, and wt was in between with a value of 3.1 px. In Fig 4D, Cdc45 was statistically significant and had a surprisingly similar pattern to Rad51 and had similar values. Cdc45 in *cds1Δ* was the longest at 4.3 px, with wt at 3.9 px, and *mrc1Δ* at 2.9 px. Interestingly, the pattern where *cds1Δ* had the longest value, wt had the mid-length, and *mrc1Δ* had the shortest length was seen in BrdU, Rad51, and Cdc45.

**Figure 4.**
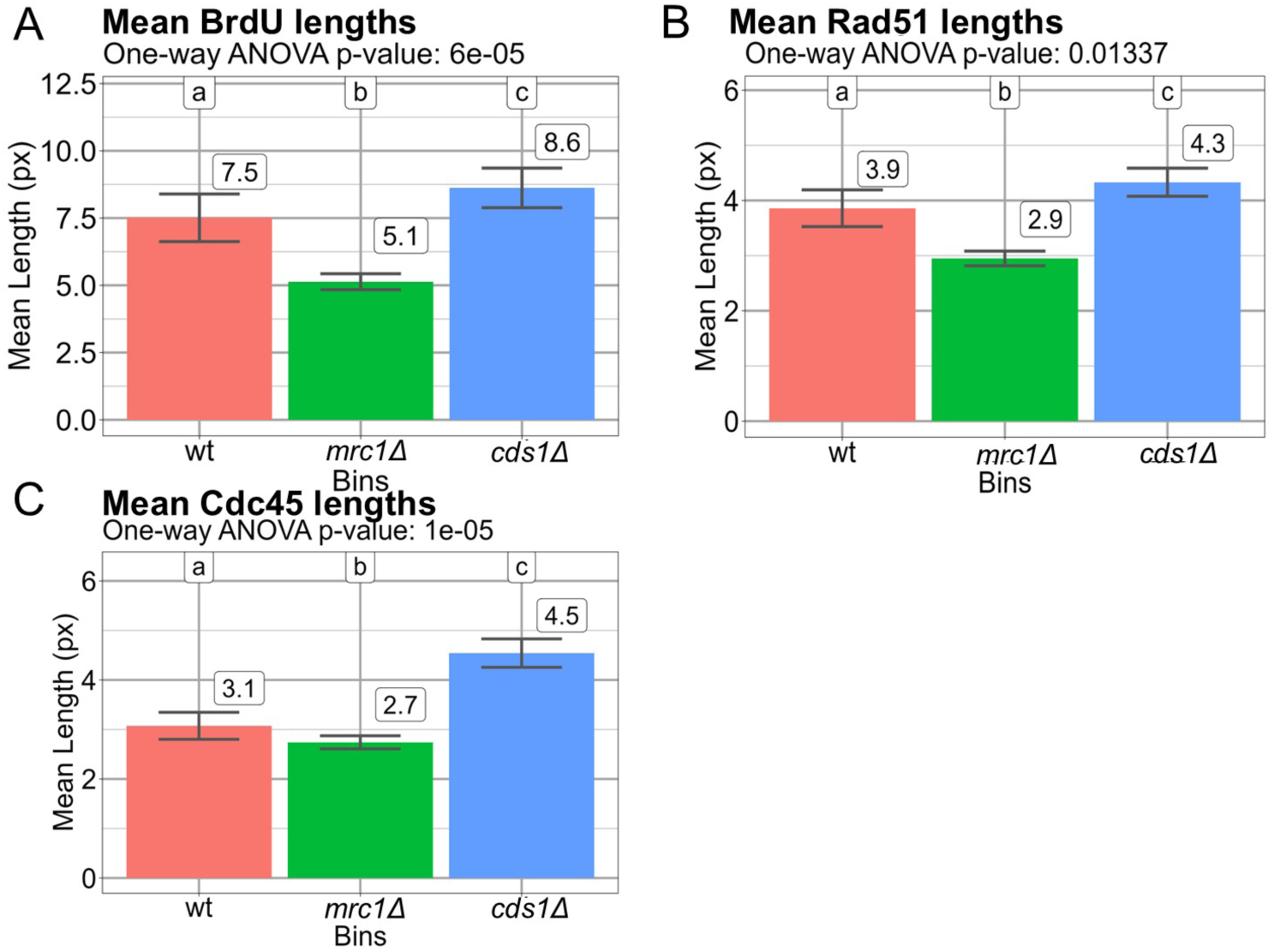
Mean lengths of BrdU, Rad51, and Cdc45 tracts calculated with optimal thresholding and smoothing parameters. Thresholding and smoothing values determined from Figures 2-3 were used to determine tract lengths of BrdU, and the proteins Rad51, and Cdc45 using R-ODD-BLOBS. Three replicates were used for each of wt, *cds1Δ*, and *mrc1Δ* cells. Cells were treated with 12 mM HU for 3.5 h, and released into fresh medium with BrdU to detect synthesis after 30 min. All values are shown in pixels (px). Variation in tract populations was compared using a 1-way ANOVA with Tukey’s HSD post-hoc correction. (A) Average BrdU-tracts post HU are longest in wt and *cds1Δ,* but shorter in *mrc1Δ.* This is consistent with previous results (Sabatinos et al 2012). There is significant variation between genotypes (p<6 × 10^−5^). (C) Rad51 recombination protein tracts are longest in *cds1Δ* tracts after HU and shortest in *mrc1Δ* (p=0.01). (D) Cdc45 lengths are similar in wt and *mrc1Δ*, and longer in *cds1Δ* (p<1 ×10^−5^).

### 3.5 Rad51 and Cdc45 co-localization at tract tips/interiors and un-replicated areas is altered in cds1Δ and mrc1Δ mutant samples

Because the smoothing function can link tracts across small gaps in signal, proteins that form long filament structures such as Rad51 can also be smoothed. We expect that smoothing protein signals will affect colocalization and tract length and is a metric that can be used to understand replication fork structures. Smoothing BrdU or protein signals may also change whether a protein is fork-associated. Because the effect of smoothing can affect individual variables (BrdU, Cdc45, Rad51 independently), or together (co-localization effects), we explored how smoothing affects observed location around tract tips. We used the previously determined smoothing baseline values (section 3.3; BrdU= 3, Rad51= 4, Cdc45= 4) and tested with no smoothing (=0 for all) to compare across individual channels. We found that protein colocalization with BrdU tracts was similar with or without smoothing in any channel (Figure 5A). BrdU smoothing alone caused a small decrease of Rad51 and Cdc45 fork co-localization in *cds1Δ* specifically, while interior tract location increased. The majority of Rad51 in Fig 5B were at unreplicated regions; *cds1Δ* had the most, and wt had the least. Around 10-28% of Rad51 were colocalized at the fork, with *mrc1Δ* having nearly double the proportion in comparison to *cds1Δ*. Colocalization at the tract was the smallest proportion at around 3-15%, where *mrc1Δ* has the least and wt and *cds1Δ* have similar proportions.

**Figure 5:**
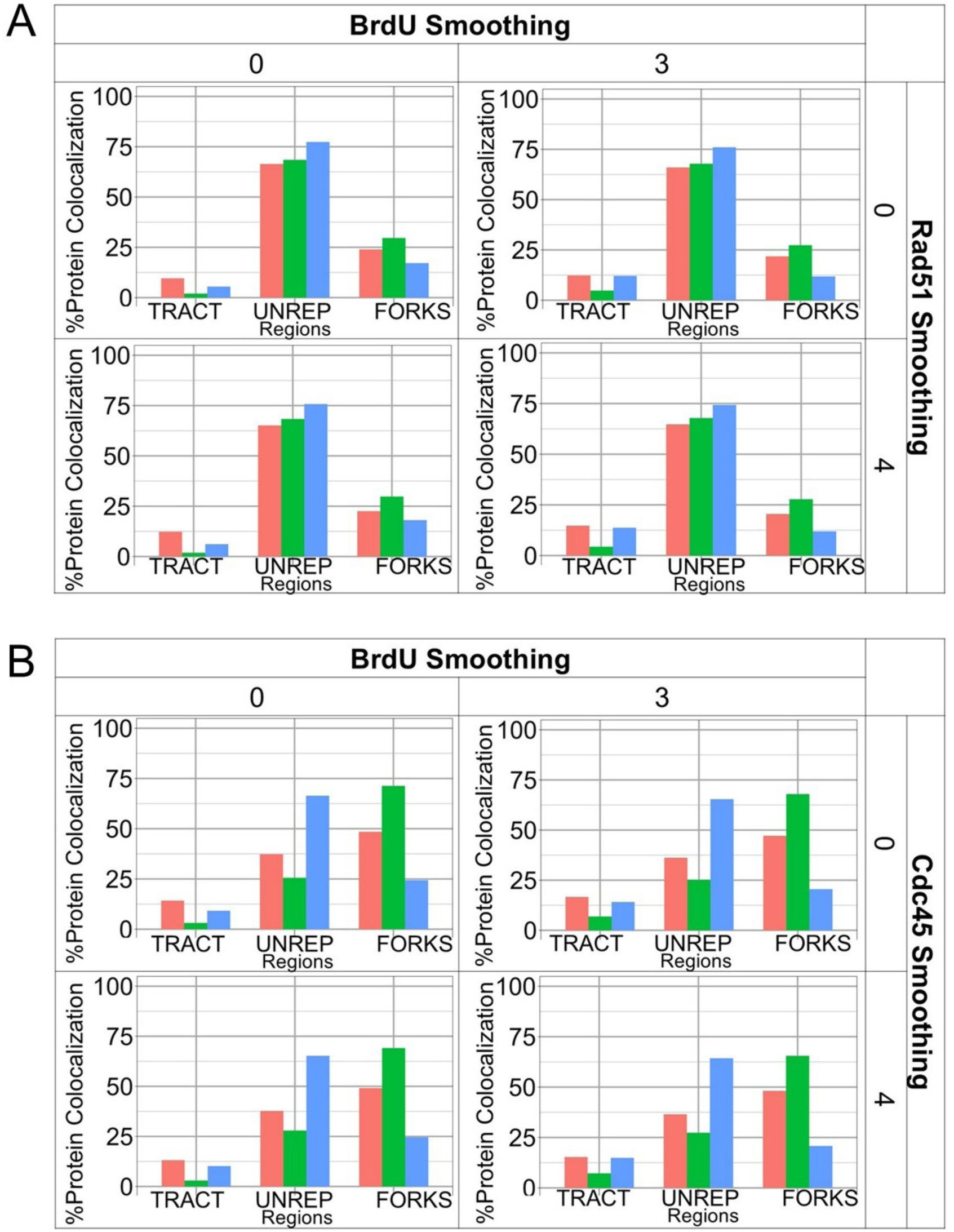
BrdU and protein smoothing parameters affect models for protein colocalization around replicated tracts. The impact of smoothing on both BrdU and/or protein (Rad51, Cdc45) was compared to each other. Full iterations of smoothing parameters are shown in Supplementary Figures. Smoothing of 0 px or 3 px was used for BrdU. Based on initial modeling, smoothing of 0 px or 4 px was compared for both Rad51 and Cdc45. In both images, wildtype is pink, *mrc1Δ* is green, and *cds1Δ* is blue. (A) Comparison of no-smoothing and smoothing on Rad51 distribution around replicated tract ends. BrdU-replicated tract tips are “FORKS”, and association with other BrdU areas is “TRACT”. Non-BrdU associated Rad51 is “UNREP” in the unreplicated areas, predicted to be remote from replication forks. (B) As in A, comparison of no-smoothing and smoothing on Cdc45 distribution around replicated tract ends.

**Figure 6.**
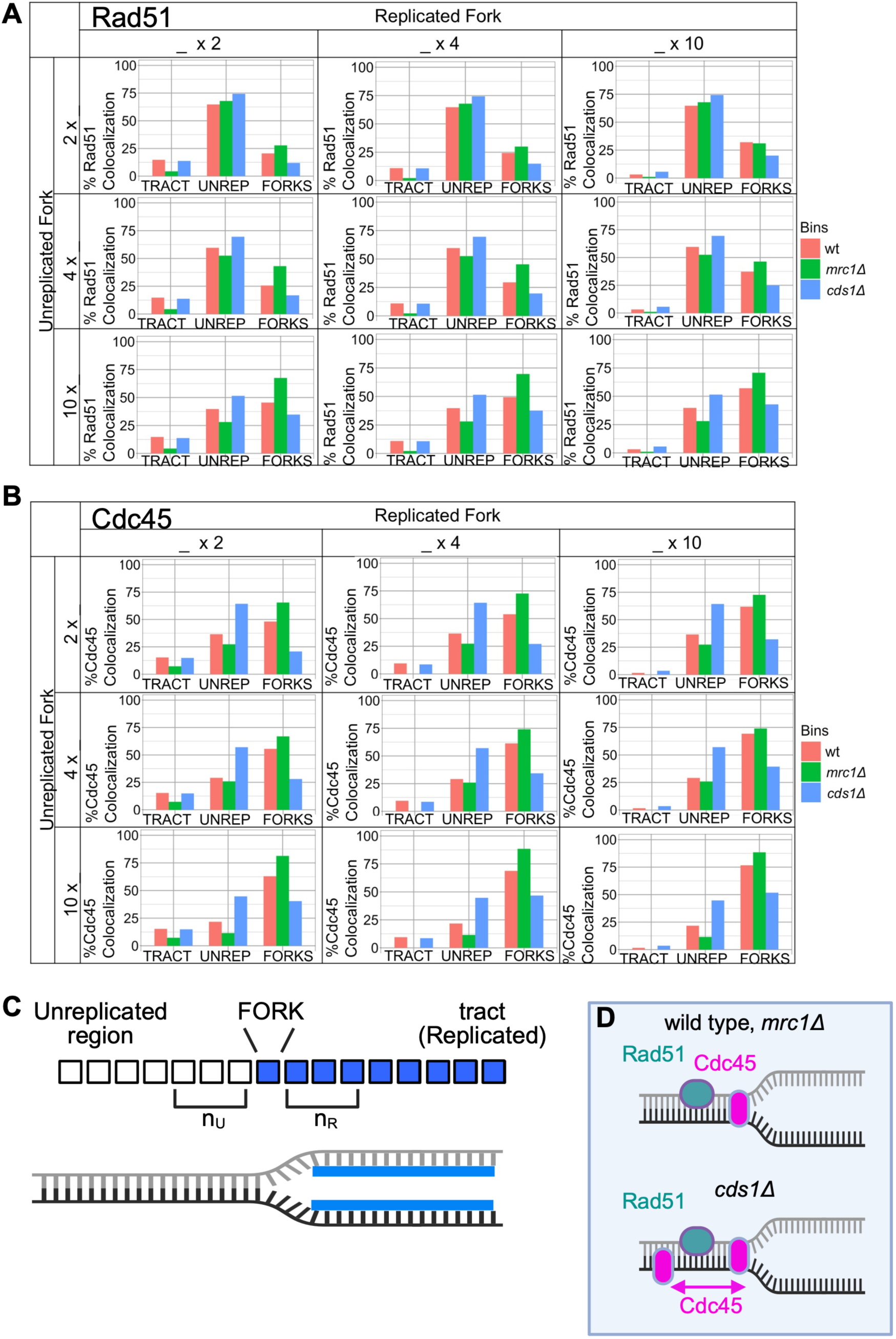
Changing the number of replicated and unreplicated fork pixels around the end of a replicated tract influences Rad51 patterns. A) Bar graphs compare iterated pixel numbers around tips relative to Rad51-region colocalization (related to Supplementary Figure S8). With no pixels included on either side of the tip, Rad51 is primarily unreplicated (“COLO UNR REGION”) regardless of fork window around BrdU tips. As the grid goes from top to bottom, the number of pixels extending in the unreplicated area near the fork increases from 2 to 10; full iteration in steps of 2 pixels is shown in Supplementary Figure S8. Rad51 located with forks increases in most in *mrc1Δ*, suggesting that *mrc1Δ* forks accumulate Rad51 that spreads from the fork into unreplicated areas. In contrast, *cds1Δ* forks retain the most Rad51 in unreplicated areas that are away from forks, suggesting DNA damage and more spread away from collapsing forks. From left to right, pixels into the replicated fork area increase from 2 to 10. Extending the number of fork-pixels into replicated area has a small effect on Rad51 localization, regardless of genotype. The 10×10 extreme situation (bottom right) is dominated by the effect of un-replicated pixels made fork-proximal. B) Analysis of Cdc45 distribution around replicated tract tips using the methos in 6B (full iterative analysis in increments of 2px is found in Supplementary Figure S8). Cdc45 is mostly un-replicated in *cds1Δ*, and is mostly tip-located in wild type and *mrc1Δ*. C) The effect of modeling pixel numbers around a tract tip is shown. Each box represents replicated (blue) or unreplicated (clear) pixels after thresholding and smoothing. The last replicated pixel “tip” is where a fork may be found. By iterating the number of pixels included as part of the fork in either the unreplicated (n_U_) or replicated (n_R_) areas, R-ODD-BLOBS can test protein co-localization patterns. In wild type cells, we predict that most Cdc45 will be close to the tip of the replicated tract; Cdc45 may move into un-replicated areas in *cds1Δ* and *mrc1Δ* preceding fork collapse. Wild type cells are expected to have Rad51 in small tracts that may be in un-replicated areas following DNA damage; in *cds1Δ* and *mrc1Δ,* Rad51 is expected to accumulate and be in un-replicated areas away from the fork. D) Summary of fork modeling for Rad51 and Cdc45 in wild type, *mrc1Δ,* and *cds1Δ* cells after HU treatment and 30 min recovery. We found that Rad51 in all strains is mostly unreplicated in all samples if there is no tip window, but unreplicated Rad51 becomes fork associated with increased pixels around tract tips.

We applied the same principles to Cdc45 distribution around replicated tips. Cdc45 distribution had a wide range from 20-65% as smoothing was increased. Unreplicated areas had most Cdc45 in *cds1Δ* (Figure 5B), and as smoothing increased the amount of Cdc45 at *cds1Δ* forks also increased. Wild type and *mrc1Δ* had similar Cdc45 distributions, arguing that Cdc45 is not completely freed from the majority of *mrc1Δ* replisomes at 30 min post HU. All genotypes and smoothing value conditions show low tract/replicated colocalization of Cdc45. We infer that any “chicken foot” structures that may form do not involve Cdc45 regression into replicated areas.

### 3.6 Fork window size explores Rad51 functions at replication forks

A key feature of R-ODD-BLOBS is that users can define the size of a fork in pixels, and rigorously test how close a protein is to the replicated tract tip. Rad51 is a recombination protein that has been immune-precipitated with replication fork components. We found that a setting the number of replicated and unreplicated pixels around the fork to 0 increased the detection of Rad51 in unreplicated areas (Figure 6A, 6C, S8). However, Rad51 filament formation in homologous recombination (HR) could extend from a broken fork into unreplicated chromatin. We increased the number of fork-associated pixels into non-BrdU areas and found that all forks have more associated Rad51 upstream. But, the increased Rad51 that was BrdU-adjacent was particularly different in *mrc1Δ* samples, suggesting that *mrc1Δ* forks are particularly sensitive to HR-restart. We thenpredicted that Rad51 might accumulate in replicated areas during HR-mediated restart post-HU. We iterated the number of replicated pixels included in the fork window (Figure 6A, 6C, S8A), but the increase in fork-association was small. We concluded that Rad51 is primarily fork associated that spreads into unreplicated DNA during HU restart, and much less into previously replicated parts of tracts (Figure 6D).

Cdc45 is a part of the CMG complex that unwinds the DNA. Both *cds1Δ* and *mrc1Δ* cells accumulate large amounts of single stranded DNA in HU arrest which we has been associated with continued stop CMG unwinding even as the polymerases are stalled. Because we might detect Cdc45 placed differently around replicated “fork” tips in different genotypes after hydroxyurea treatment, we tested how changing the fork window altered Cdc45 location (Figure 6B, S8B). Increasing the number of unreplicated or replicated pixels at a tip had small effects on wild type and *mrc1Δ* becauseCdc45 is mostly at the tract tip/fork. In contrast, *cds1Δ* cells showed most Cdc45 at unreplicated areas, regardless of window size. It was only at the extreme of 10 px (∼3 kb of DNA) added around tips that *cds1Δ* Cdc45 became more tip-associated. We concluded that Cdc45 is fork-associated in wild type and *mrc1Δ* cells (Figure 6D), but is spread away from forks into un-replicated areas in *cds1Δ* samples during HU recovery.

## 4.0 CONCLUSIONS

### 4.1 R-ODD-BLOBS provides opportunities to iterate and understand replication fork biology

Our work shows that high-resolution analysis of chromatin fiber data can model replication protein location and structures. Because of our preparation method that fixes samples in time, we predict that forks will be located at the end of the replicated tracts. We have shown that the ability to test threshold values, smoothing parameters, and fork window sizes provide important clues to variables that define replication forks. Our R-based code has improved the stability and speed of R-ODD-BLOBS analysis, allowing iterative analysis. We found optimal thresholding parameters for these data which allowed fork-window modeling (Figure 6), leading to a conclusion that Cdc45 spreads into unreplicated areas away from *cds1Δ* forks. Thus, *cds1Δ* forks accumulate single stranded DNA because CMG is uncoupled or is dissociated from replicating forks. Rad51 functions in homologous recombination restart at forks, consistent with R-ODD-BLOBS modeling that shows Rad51 is at forks and tends to spread into unreplicated areas if the fork window is sufficiently large.

Our analysis method is faster than manual analysis. Co-localization and measurement uses defined thresholding, smoothing, and tip window parameters. These provide a rigorous framework that can be used to link many DNA replication fork proteins from multiple datasets. R-ODD-BLOBS facilitates individual or binned graphs of protein-region colocalization from datasets (*i.e*. Figures 5 and 6). Binning brings together multiple experimental replicate observations. Because colocalization data are categorical, binning within populations indicates trend consistency and size for comparison between strain conditions. We also included features to bin lengths for each channel over multiple datasets (seen in Figure 4), or to test the length of each channel compared to each dataset and initial scatter plots (Supplementary Figure S1). These graphs can help users quickly identify patterns within their data and to facilitate data analysis.

While thresholding for intensity has easily predictable effects on BrdU or protein tracts, smoothing tests whether gaps in signal are real or insignificant. The Abbé limit guides smoothing for signal gaps below the limit of light resolution. Using a green light excitation (488 nm), NA and pixel size for the system (1.4 NA, 0.1092 µm/px), the Abbé limit states that gaps in BrdU-signal less than 3 pixels long cannot be discriminated. Standard smoothing values of 3px for green (BrdU), and 4px for orange (Rad51, 546 nM) and red (Cdc45, 647 nm) were tested and do not affect the patterns observed. However, care must be taken with smoothing, since large values of smoothing will connect gaps into longer lengths (Figure 3, Supplmentary Figures S5-S7).

### 4.2 Building replication fork models from averaged datasets

There are significant differences between the mean lengths of BrdU in wt, *cds1Δ* and *mrc1Δ*, as seen in Fig 4B. *mrc1Δ* is the shortest at 5.1 px, *cds1Δ* is the longest at 8.6 px, and wt is right behind at 7.5 px. This trend is comparable to literature where *cds1Δ* was the longest, followed by wt and then *mrc1Δ* which was less than half of *cds1Δ* [23] after a 45 min release from HU. We propose that *mrc1Δ* synthesis is most impacted, despite Cdc45 proximity to fork areas.

Cdc45 is a part of the CMG complex with with Mcm2-7 and GINS that unwinds dsDNA during replication [43]. We found that Cdc45 is away from forks in *cds1Δ,* indicating that replication forks activities of synthesis and fail to recover by 30 min after release leading to truncated replication [23]. Cdc45 could appear at unreplicated regions during origin firing [44] or after pre-replication complexes are destroyed [45]. Increased unreplicated Cdc45 in *cds1Δ* could represent uncoupling between helicase and polymerase subunits, or that Cds1 regulates origin association leading to improper Cdc45 binding post-HU. While Cdc45 in both *mrc1Δ* and wild type are similar and fork-proximal, our next work will calculate tract length and fluorescent intensity to model the differences between wildtype, which survives HU, and *mrc1Δ,* which dies in HU.

The structure of any given fork and its associated proteins is loosely defined, particularly in how protein loss leads to fork collapse and the generation of DNA damage. However, proteins such as Rad51 can filament during homolgous recombination detection and decoration of single stranded DNA. Without increasing fork inclusion on either side of a tip, Rad51 might be recognized different protein events that are beside each other. Our analysis examines increasing pixels away from a “fork window” on either side of the replicated tip “FORK”. We predicted that R-ODD-BLOBS can identify whether Rad51 is spread between regions or connected to the fork region. Similarities between *cds1Δ* and wt Rad51 protein colocalization suggest that lethal levels of DNA damage have not occurred within 30 minutes after the release of HU, and that de-regulated *cds1Δ* late origin firing, and DNA unwinding [23] coincident with Cdc45 patterns might promote *cds1Δ* death that is realized later in growing colonies.

R-ODD-BLOBS provides a unique opportunity to understand replication fork size and spatial factors. For example, Rad51 assembles into a filament on RPA to coat ssDNA during HR [24]; our data in Fig 6 reflects this and shows linked Rad51 colocalization to the fork and unreplicated regions suggesting filaments. Future work using Rad52 (a mediator of Rad51 in yeast [26]) can determine if the Rad51 spread is from fork restart or recombination repair. This iterative nature give R-ODD-BLOBS the ability to model *cds1Δ, mrc1Δ,* and other DNA replication instability mutants, while deconvolving epigenetic and aggregate evidence for the similar phenotypes but different roles of Cds1 and Mrc1 [40,46,47]. Our next steps will correlate single-fork architectures with neighboring fork complexes to determine the variety of fork patterns across a genome and relative to each other in a chromatin context. We also recognize the power of 2-color labeling (with IdU and CldU), and specific genomic areas detected by FISH (i.e Fig 1), to model DNA replication instability.

## Supporting information

All supplementary figures

## Acknowledgements

We thank JiPing Yuan and Marc Green (USC) for their work and consultation in the development of this project. We thank Uzair Mayat and Aniqa Imtiaz important discussions on code-conversion and spatial applications to SynLab activities. This work was begun with support from NIH R01 GM059321 to SLF. This work was funded by the Canadian Natural Science and Engineering Research Council (NSERC) to SAS (RGPIN-2015-04405) and AM (Discovery Grant Program), in addition TMU Faculty of Science NSERC Booster and Bridge support to SAS, NSERC USRA awards to KA, and TMU URO funding to KA.

## ABBREVIATIONS

DSB: Double strand break
BrdU: bromodeoxyuridine
*S. pombe*: *Schizosaccharomyces pombe*
HR: homologous recombination
dsDNA: double, strand DNA
ssDNA: single stranded DNA
ODD-BLOBS: One Dimensional Data – Boolean Logic Operations Binning System

## 7.0 TABLES

| Genotype | Source |
| --- | --- |
| *h<sup>+</sup> rad11myc::KanMX cdc45-YFP-ura4<sup>+</sup> leu1-32::hENT1-leu1+(pJAH29) his7-366::hsv-tk-his7+(pJAH31) ura4-D18 ade6-M210* | FY 3805* |
| *h<sup>+</sup> Δmrc1::ura4<sup>+</sup> cdc45-YFP-ura4<sup>+</sup> leu1-32::hENT1-leu1+(pJAH29) his7-366::hsv-tk-his7+(pJAH31) rad11myc::KanMX ura4-D18 ade6-M210* | FY 3821 |
| *h<sup>+</sup> Δcds1::ura4+ cdc45-YFP-ura4+ rad11myc::KanMX leu1-32::hENT1-leu1+(pJAH29) his7-366::hsv-tk-his7+(pJAH31) ura4-D18 ade6-M210 (rad11=ssb1)* | FY 3830 |
| *h<sup>-</sup> pola-FLAG2::ura4+ rad11myc::KanMX leu1-32::hENT1-leu1+(pJAH29) his7-366::hsv-tk-his7+(pJAH31) ura4-D18* | FY 3858 |
| *h<sup>-</sup> Δmrc1::ura4+ pola-FLAG2::ura4+ rad11myc::KanMX leu1-32::hENT1-leu1+(pJAH29) his7-366::hsv-tk-his7+(pJAH31) ura4-D18* | FY 3859 |
| *h<sup>-</sup> Δcds1::ura4+ pola-FLAG2::ura4+ rad11myc::KanMX leu1-32::hENT1-leu1+(pJAH29) his7-366::hsv-tk-his7+(pJAH31) ura4-D18* | FY 3866 |

\*Forsburg Yeast (FY) collection, SL Forsburg, University of Southern California

## 8.1 Supplementary Figure legends

**Supplemental Figure 1:**
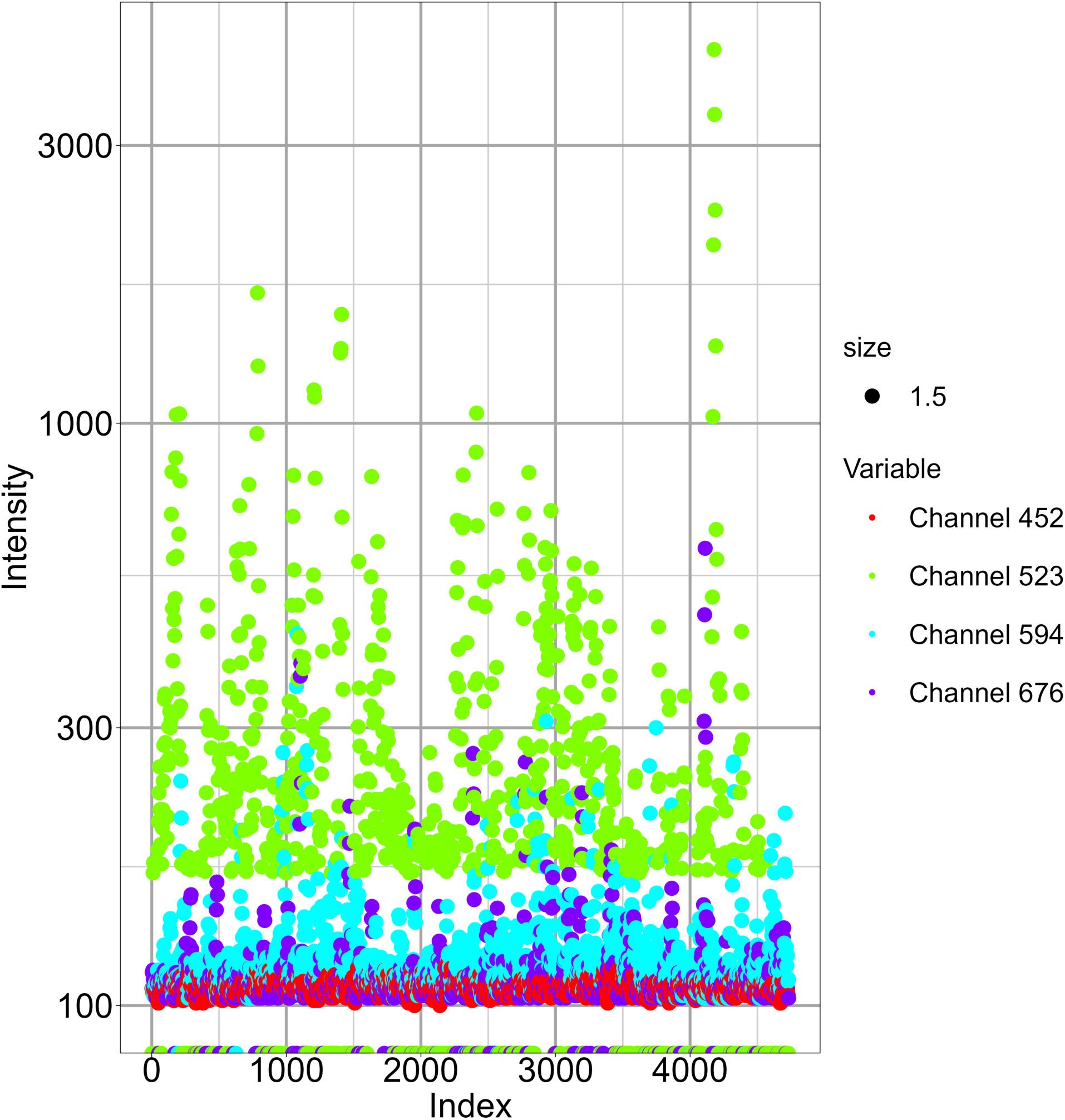
Initial Scatter Plot of Channel intensities from a single wt image.

**Supplemental Figure 2:**
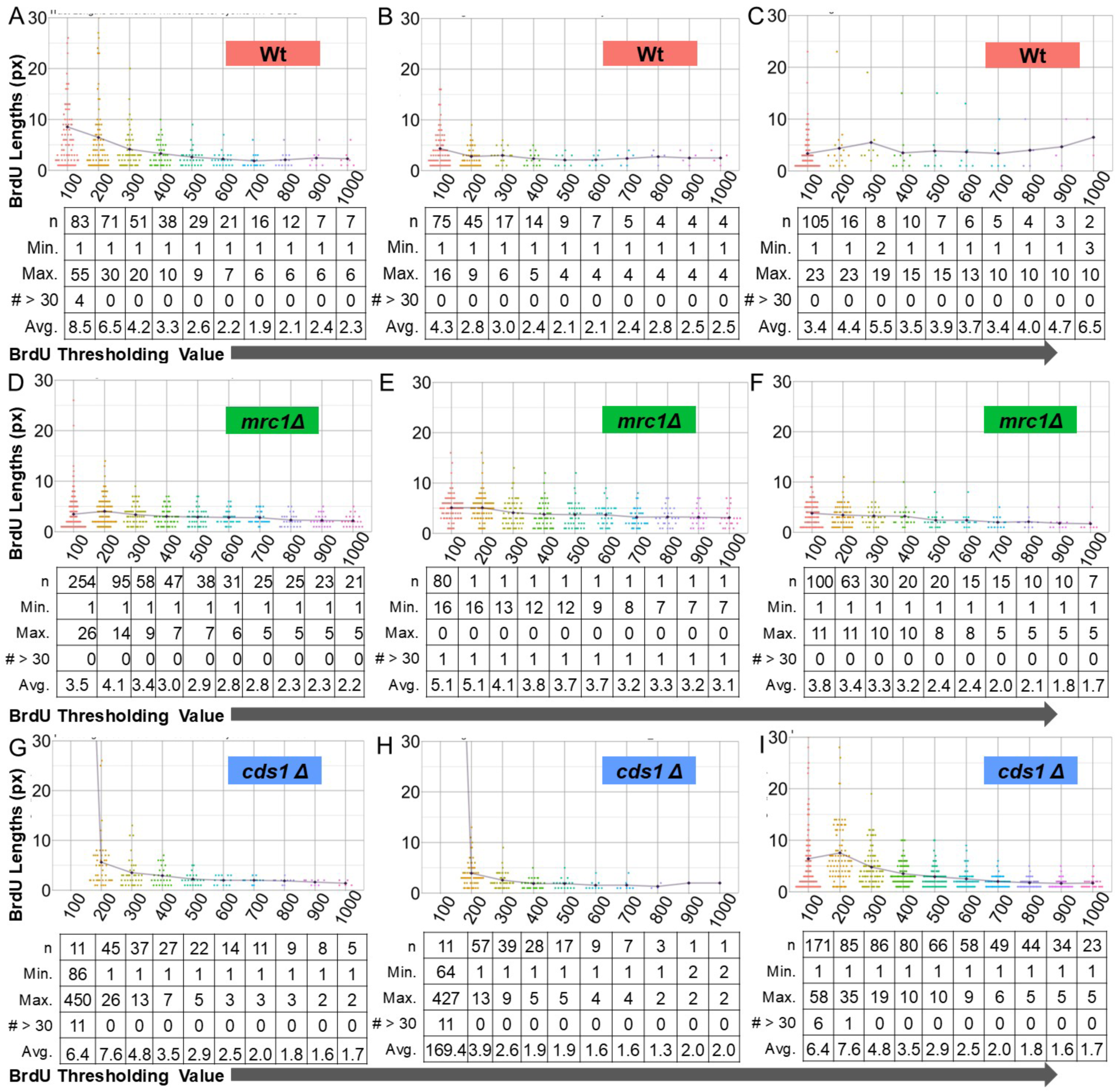
Individual Replicates of BrdU Thresholding for Wt, *cds1Δ* and *mrc1Δ*.

**Supplemental Figure 3:**
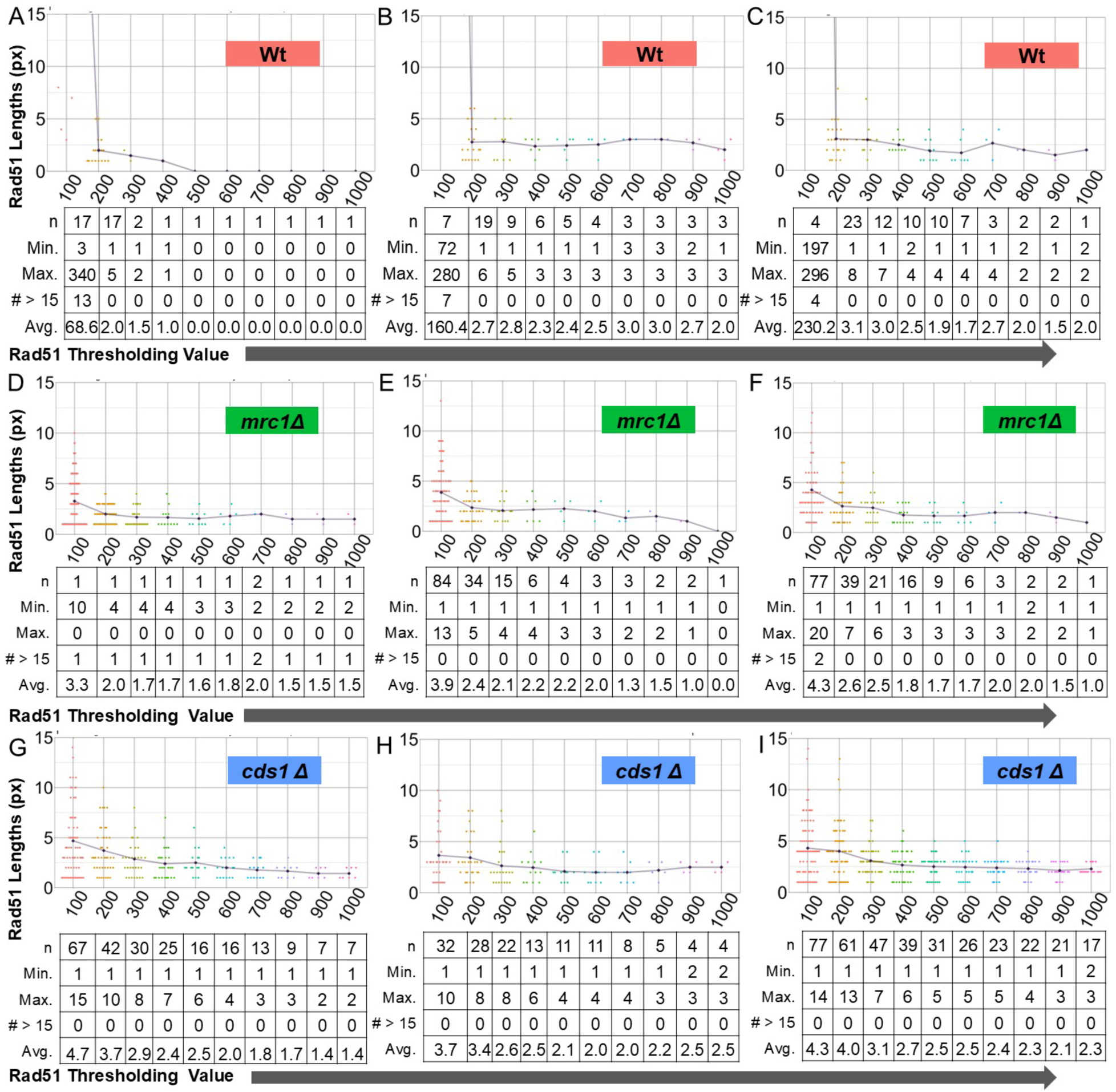
Individual Replicates of Rad51 Thresholding for Wt, *cds1Δ* and *mrc1Δ*.

**Supplemental Figure 4:**
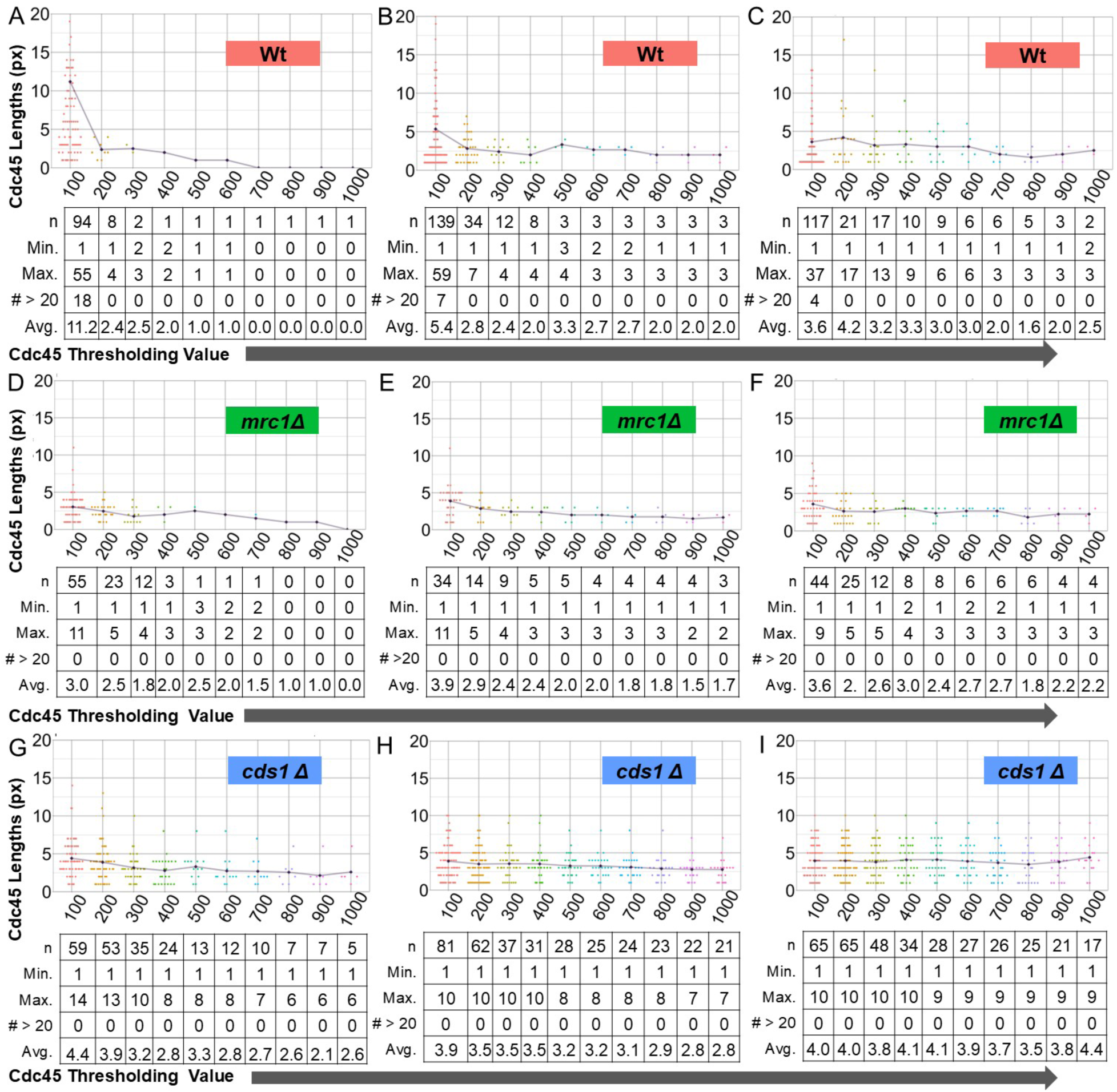
Individual Replicates of Cdc45 Thresholding for Wt, *cds1Δ* and *mrc1Δ*.

**Supplemental Figure 5:**
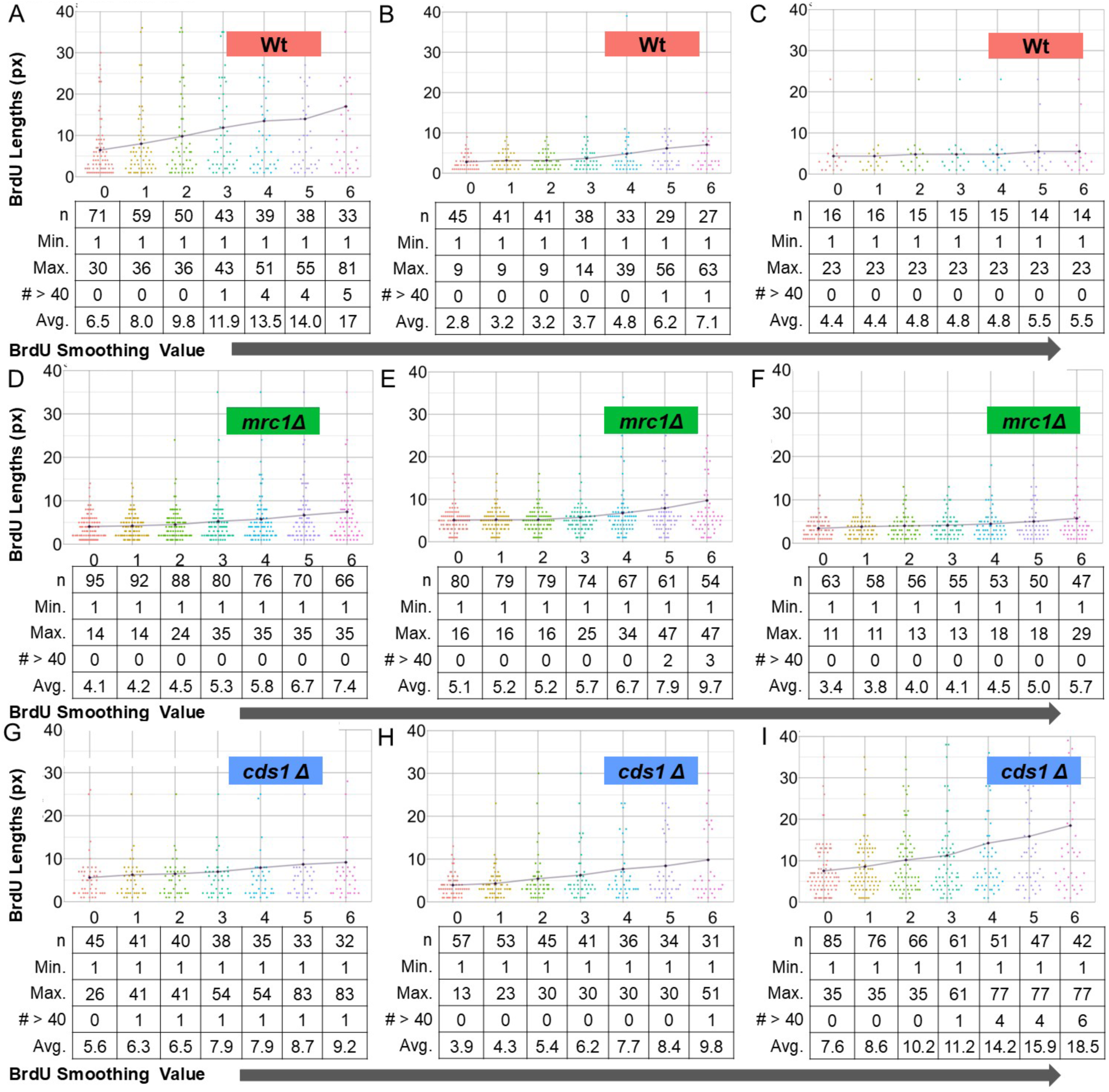
Individual Replicates of BrdU Smoothing for Wt, *cds1Δ* and *mrc1Δ*.

**Supplemental Figure 6:**
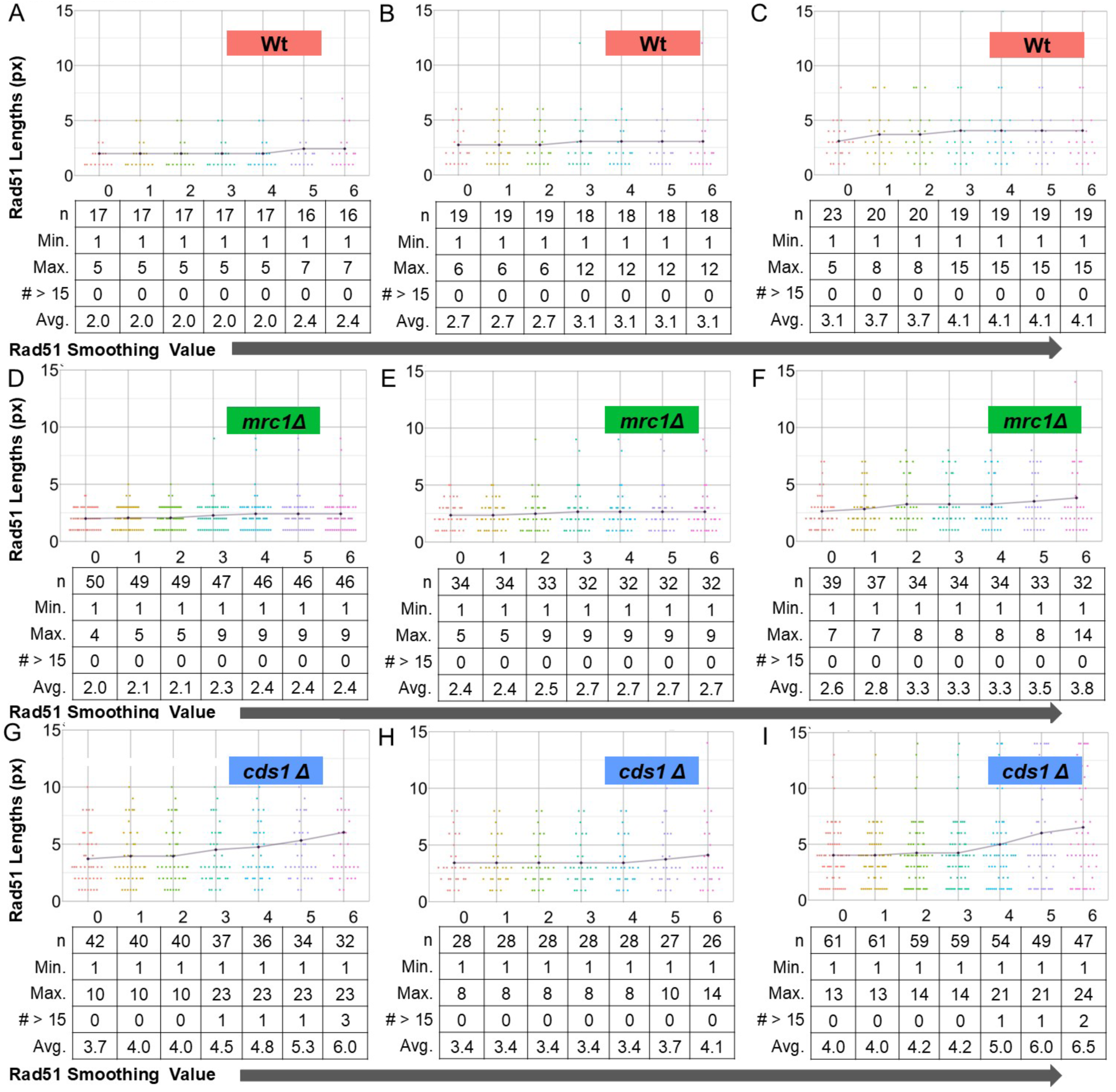
Individual Replicates of Rad51 Smoothing for Wt, *cds1Δ* and *mrc1Δ*.

**Supplemental Figure 7:**
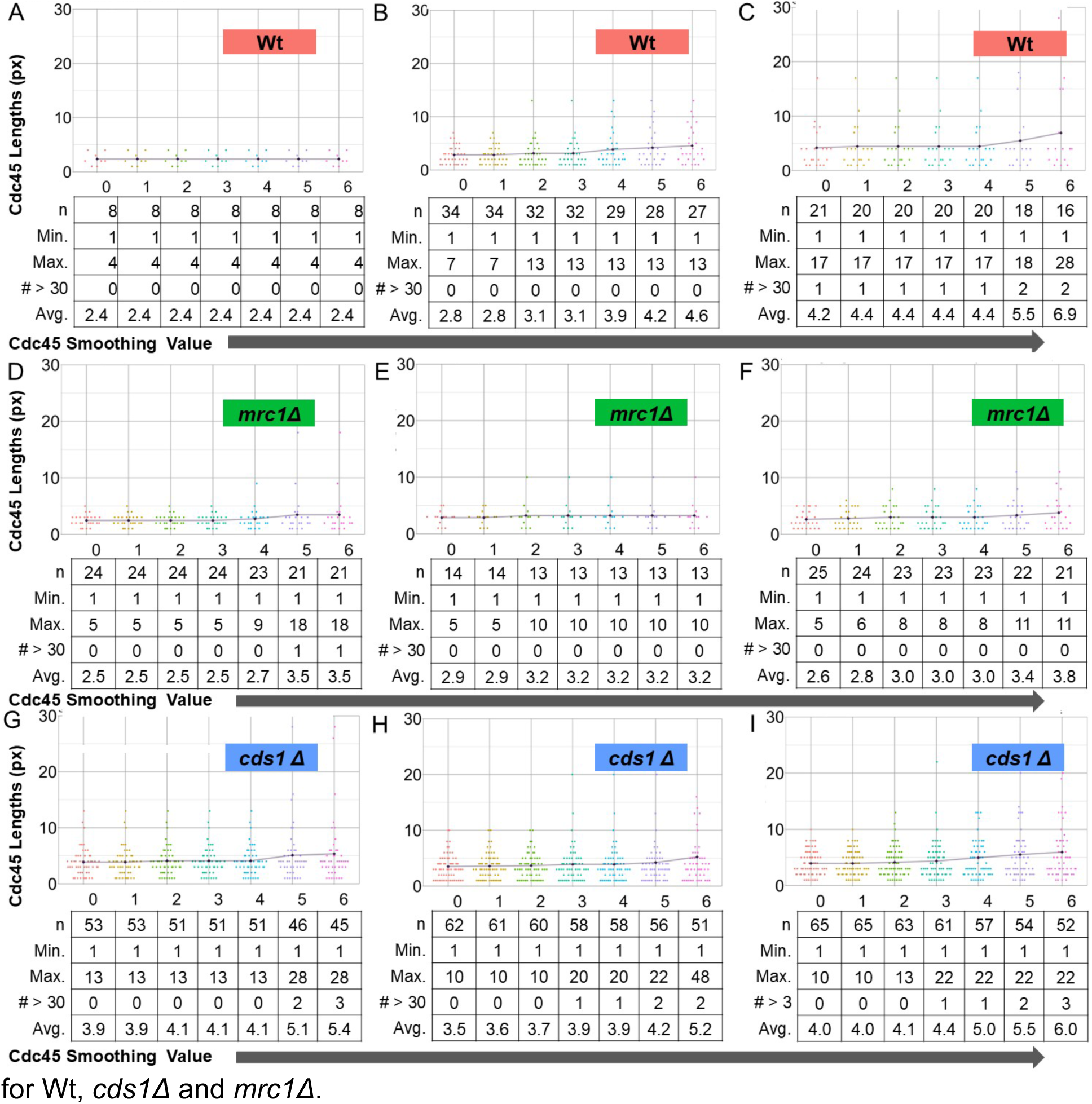
Individual Replicates of Cdc45 Smoothing iteration. for Wt, *cds1Δ* and *mrc1Δ*.

**Supplemental Figure 8: The effect of iterative window sizing on Rad51 and Cdc45 distribution near forks.** A) Rad51 located with forks increases in most in *mrc1Δ*, suggesting that *mrc1Δ* forks accumulate Rad51 that spreads from the fork into unreplicated areas. In contrast, *cds1Δ* forks retain the most Rad51 in unreplicated areas that are away from forks, suggesting DNA damage and more spread away from collapsing forks. From left to right, pixels into the replicated fork area increase from 2 to 10. Extending the number of fork-pixels into replicated area has a small effect on Rad51 localization, regardless of genotype. The 10×10 extreme situation (bottom right) is dominated by the effect of un-replicated pixels made fork-proximal.

B) Expanded window-size effects on Cdc45 colocalization for *cds1Δ (pink), mrc1Δ (green),* and wild type (blue). With no pixels included on either side of the tip, Cdc45 is primarily unreplicated (“COLO UNR REGION”) regardless of fork window around BrdU tips. As the grid goes from top to bottom, the number of pixels extending in the unreplicated area near the fork increases from 2 to 10.

