## Supplementary material for "Modelling DNA replication fork stability and collapse using chromatin fiber analysis and the R-ODD-BLOBS program": All supplementary figures

***SUPPLEMENTARY INFORMATION***

**AUTHORS**:
Kerenza Cheng[1], Kazeera Aliar[2], Roozbeh Manshaei[3], Susan L Forsburg[4], Ali Mazalek[1,3], Sarah A Sabatinos[1,2, *]

**AFFILIATIONS**

1 – Molecular Science Graduate Program, Yeates School of Graduate and Postdoctoral Studies, Toronto Metropolitan University, Toronto ON M5B 2K3

2 – Department of Chemistry and Biology, Toronto Metropolitan University, Toronto ON M5B 2K3

3 – Synaesthetic Media Lab, The Creative School, Toronto Metropolitan University, Toronto ON M5B 2K3

4 – Department of Molecular and Computational Biology, University of Southern California, Los Angeles, California USA 90089

**Supplemental Figure 1:** Initial Scatter Plot of Channel intensities of One WT file
**Supplemental Figure 2:** Individual Replicates of BrdU Thresholding for Wt, *cds1Δ* and *mrc1Δ*.
**Supplemental Figure 3:** Individual Replicates of Rad51 Thresholding for Wt, *cds1Δ* and *mrc1Δ*.
**Supplemental Figure 4:** Individual Replicates of Cdc45 Thresholding for Wt, *cds1Δ* and *mrc1Δ*.
**Supplemental Figure 5:** Individual Replicates of BrdU Smoothing for Wt, *cds1Δ* and *mrc1Δ*.
**Supplemental Figure 6:** Individual Replicates of Rad51 Smoothing for Wt, *cds1Δ* and *mrc1Δ*.
**Supplemental Figure 7:** Individual Replicates of Cdc45 Smoothing for Wt, *cds1Δ* and *mrc1Δ*.
**Supplemental Figure S8:** Effects of Replication Fork Window size on Rad51 and Cdc45 colocalization

**Supplemental Figure 1:** Initial Scatter Plot of Channel intensities from a single wt image.

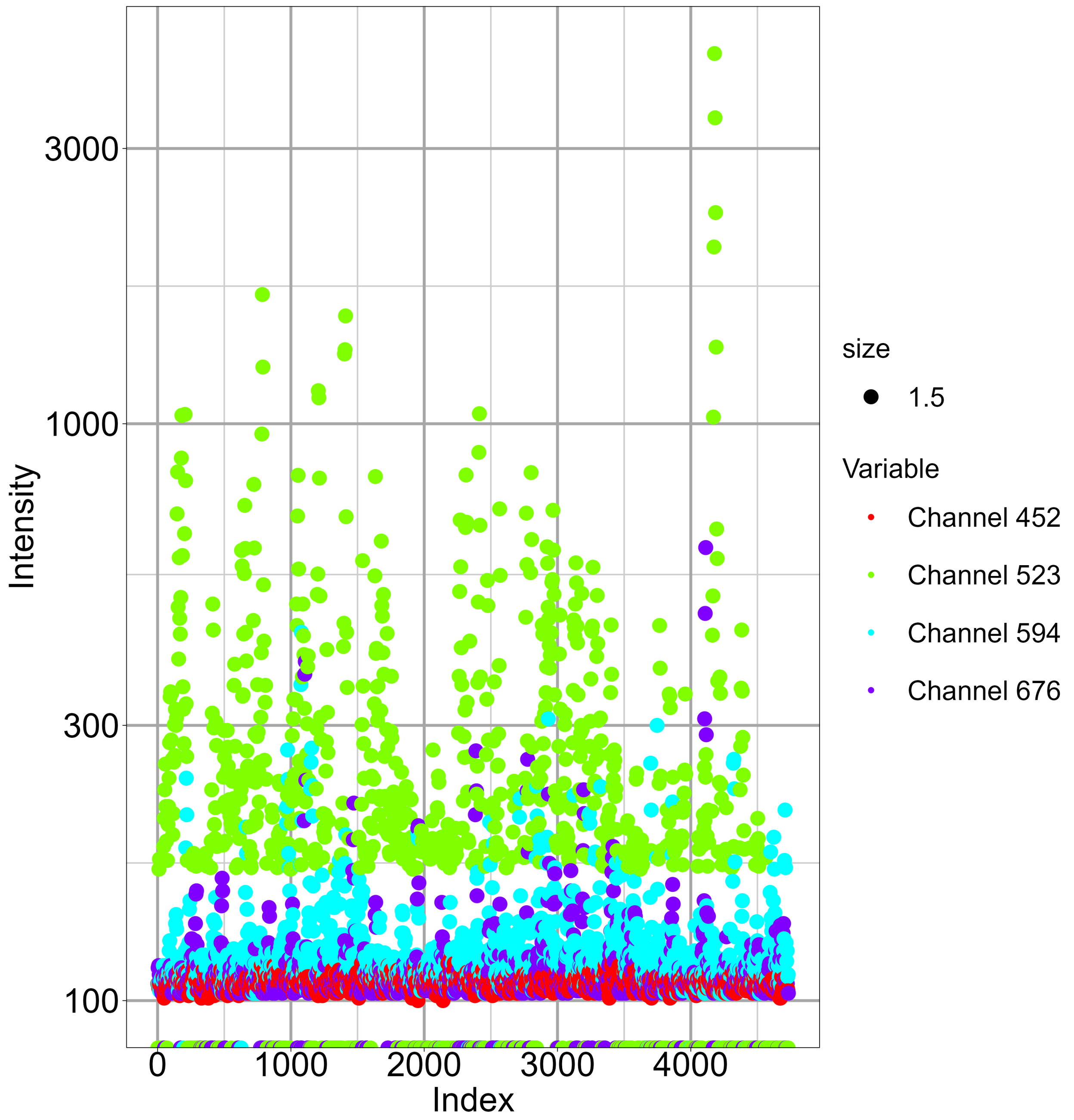

**Supplemental Figure 2:** Individual Replicates of BrdU Thresholding for Wt, *cds1Δ* and *mrc1Δ*.

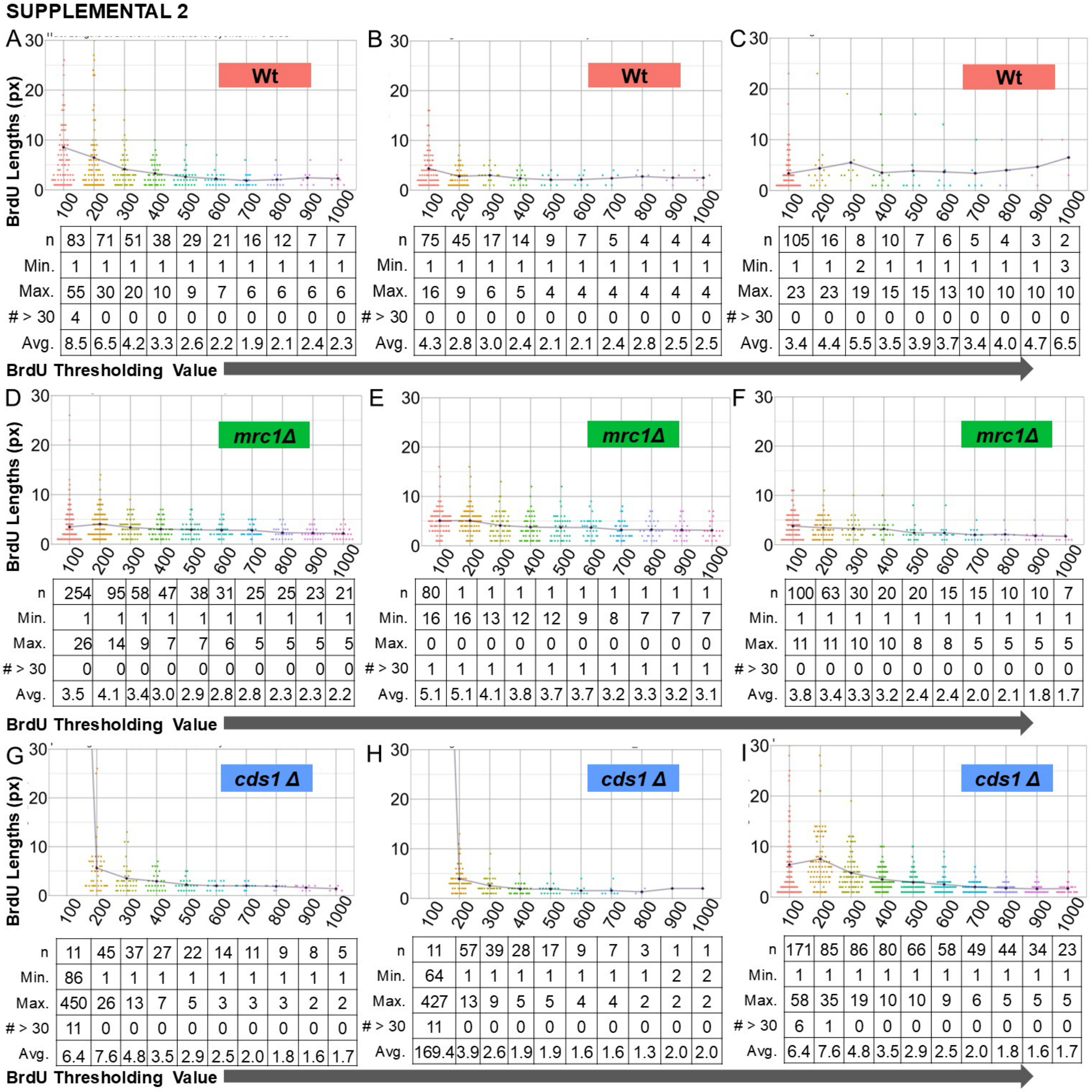


A (Wt)

| | 100 | 200 | 300 | 400 | 500 | 600 | 700 | 800 | 900 | 1000 |
|---|---|---|---|---|---|---|---|---|---|---|
| n | 83 | 71 | 51 | 38 | 29 | 21 | 16 | 12 | 7 | 7 |
| Min. | 1 | 1 | 1 | 1 | 1 | 1 | 1 | 1 | 1 | 1 |
| Max. | 55 | 30 | 20 | 10 | 9 | 7 | 6 | 6 | 6 | 6 |
| # > 30 | 4 | 0 | 0 | 0 | 0 | 0 | 0 | 0 | 0 | 0 |
| Avg. | 8.5 | 6.5 | 4.2 | 3.3 | 2.6 | 2.2 | 1.9 | 2.1 | 2.4 | 2.3 |

B (Wt)

| | 100 | 200 | 300 | 400 | 500 | 600 | 700 | 800 | 900 | 1000 |
|---|---|---|---|---|---|---|---|---|---|---|
| n | 75 | 45 | 17 | 14 | 9 | 7 | 5 | 4 | 4 | 4 |
| Min. | 1 | 1 | 1 | 1 | 1 | 1 | 1 | 1 | 1 | 1 |
| Max. | 16 | 9 | 6 | 5 | 4 | 4 | 4 | 4 | 4 | 4 |
| # > 30 | 0 | 0 | 0 | 0 | 0 | 0 | 0 | 0 | 0 | 0 |
| Avg. | 4.3 | 2.8 | 3.0 | 2.4 | 2.1 | 2.1 | 2.4 | 2.8 | 2.5 | 2.5 |

C (Wt)

| | 100 | 200 | 300 | 400 | 500 | 600 | 700 | 800 | 900 | 1000 |
|---|---|---|---|---|---|---|---|---|---|---|
| n | 105 | 16 | 8 | 10 | 7 | 6 | 5 | 4 | 3 | 2 |
| Min. | 1 | 1 | 2 | 1 | 1 | 1 | 1 | 1 | 1 | 3 |
| Max. | 23 | 23 | 19 | 15 | 15 | 13 | 10 | 10 | 10 | 10 |
| # > 30 | 0 | 0 | 0 | 0 | 0 | 0 | 0 | 0 | 0 | 0 |
| Avg. | 3.4 | 4.4 | 5.5 | 3.5 | 3.9 | 3.7 | 3.4 | 4.0 | 4.7 | 6.5 |

D (mrc1Δ)

| | 100 | 200 | 300 | 400 | 500 | 600 | 700 | 800 | 900 | 1000 |
|---|---|---|---|---|---|---|---|---|---|---|
| n | 254 | 95 | 58 | 47 | 38 | 31 | 25 | 25 | 23 | 21 |
| Min. | 1 | 1 | 1 | 1 | 1 | 1 | 1 | 1 | 1 | 1 |
| Max. | 26 | 14 | 9 | 7 | 7 | 6 | 5 | 5 | 5 | 5 |
| # > 30 | 0 | 0 | 0 | 0 | 0 | 0 | 0 | 0 | 0 | 0 |
| Avg. | 3.5 | 4.1 | 3.4 | 3.0 | 2.9 | 2.8 | 2.8 | 2.3 | 2.3 | 2.2 |

E (mrc1Δ)

| | 100 | 200 | 300 | 400 | 500 | 600 | 700 | 800 | 900 | 1000 |
|---|---|---|---|---|---|---|---|---|---|---|
| n | 80 | 1 | 1 | 1 | 1 | 1 | 1 | 1 | 1 | 1 |
| Min. | 16 | 16 | 13 | 12 | 12 | 9 | 8 | 7 | 7 | 7 |
| Max. | 0 | 0 | 0 | 0 | 0 | 0 | 0 | 0 | 0 | 0 |
| # > 30 | 1 | 1 | 1 | 1 | 1 | 1 | 1 | 1 | 1 | 1 |
| Avg. | 5.1 | 5.1 | 4.1 | 3.8 | 3.7 | 3.7 | 3.2 | 3.3 | 3.2 | 3.1 |

F (mrc1Δ)

| | 100 | 200 | 300 | 400 | 500 | 600 | 700 | 800 | 900 | 1000 |
|---|---|---|---|---|---|---|---|---|---|---|
| n | 100 | 63 | 30 | 20 | 20 | 15 | 15 | 10 | 10 | 7 |
| Min. | 1 | 1 | 1 | 1 | 1 | 1 | 1 | 1 | 1 | 1 |
| Max. | 11 | 11 | 10 | 10 | 8 | 8 | 5 | 5 | 5 | 5 |
| # > 30 | 0 | 0 | 0 | 0 | 0 | 0 | 0 | 0 | 0 | 0 |
| Avg. | 3.8 | 3.4 | 3.3 | 3.2 | 2.4 | 2.4 | 2.0 | 2.1 | 1.8 | 1.7 |

G (cds1 Δ)

| | 100 | 200 | 300 | 400 | 500 | 600 | 700 | 800 | 900 | 1000 |
|---|---|---|---|---|---|---|---|---|---|---|
| n | 11 | 45 | 37 | 27 | 22 | 14 | 11 | 9 | 8 | 5 |
| Min. | 86 | 1 | 1 | 1 | 1 | 1 | 1 | 1 | 1 | 1 |
| Max. | 450 | 26 | 13 | 7 | 5 | 3 | 3 | 3 | 2 | 2 |
| # > 30 | 11 | 0 | 0 | 0 | 0 | 0 | 0 | 0 | 0 | 0 |
| Avg. | 6.4 | 7.6 | 4.8 | 3.5 | 2.9 | 2.5 | 2.0 | 1.8 | 1.6 | 1.7 |

H (cds1 Δ)

| | 100 | 200 | 300 | 400 | 500 | 600 | 700 | 800 | 900 | 1000 |
|---|---|---|---|---|---|---|---|---|---|---|
| n | 11 | 57 | 39 | 28 | 17 | 9 | 7 | 3 | 1 | 1 |
| Min. | 64 | 1 | 1 | 1 | 1 | 1 | 1 | 1 | 2 | 2 |
| Max. | 427 | 13 | 9 | 5 | 5 | 4 | 4 | 2 | 2 | 2 |
| # > 30 | 11 | 0 | 0 | 0 | 0 | 0 | 0 | 0 | 0 | 0 |
| Avg. | 169.4 | 3.9 | 2.6 | 1.9 | 1.9 | 1.6 | 1.6 | 1.3 | 2.0 | 2.0 |

I (cds1 Δ)

| | 100 | 200 | 300 | 400 | 500 | 600 | 700 | 800 | 900 | 1000 |
|---|---|---|---|---|---|---|---|---|---|---|
| n | 171 | 85 | 86 | 80 | 66 | 58 | 49 | 44 | 34 | 23 |
| Min. | 1 | 1 | 1 | 1 | 1 | 1 | 1 | 1 | 1 | 1 |
| Max. | 58 | 35 | 19 | 10 | 10 | 9 | 6 | 5 | 5 | 5 |
| # > 30 | 6 | 1 | 0 | 0 | 0 | 0 | 0 | 0 | 0 | 0 |
| Avg. | 6.4 | 7.6 | 4.8 | 3.5 | 2.9 | 2.5 | 2.0 | 1.8 | 1.6 | 1.7 |

**Supplemental Figure 3:** Individual Replicates of Rad51 Thresholding for Wt, *cds1Δ* and *mrc1Δ*.

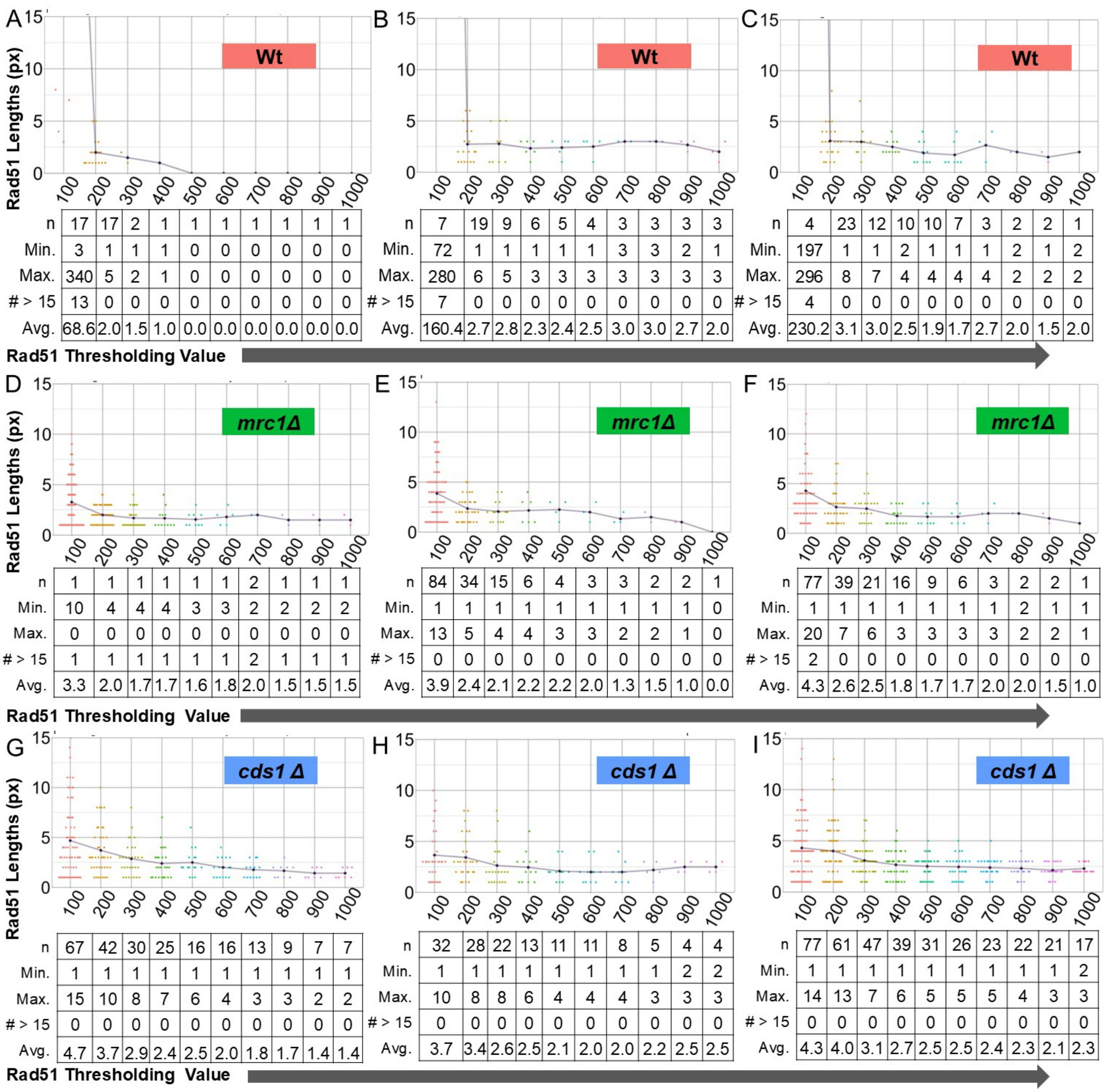


**A (Wt)**

| | 100 | 200 | 300 | 400 | 500 | 600 | 700 | 800 | 900 | 1000 |
|---|---|---|---|---|---|---|---|---|---|---|
| n | 17 | 17 | 2 | 1 | 1 | 1 | 1 | 1 | 1 | 1 |
| Min. | 3 | 1 | 1 | 1 | 0 | 0 | 0 | 0 | 0 | 0 |
| Max. | 340 | 5 | 2 | 1 | 0 | 0 | 0 | 0 | 0 | 0 |
| # > 15 | 13 | 0 | 0 | 0 | 0 | 0 | 0 | 0 | 0 | 0 |
| Avg. | 68.6 | 2.0 | 1.5 | 1.0 | 0.0 | 0.0 | 0.0 | 0.0 | 0.0 | 0.0 |

**B (Wt)**

| | 100 | 200 | 300 | 400 | 500 | 600 | 700 | 800 | 900 | 1000 |
|---|---|---|---|---|---|---|---|---|---|---|
| n | 7 | 19 | 9 | 6 | 5 | 4 | 3 | 3 | 3 | 3 |
| Min. | 72 | 1 | 1 | 1 | 1 | 1 | 3 | 3 | 2 | 1 |
| Max. | 280 | 6 | 5 | 3 | 3 | 3 | 3 | 3 | 3 | 3 |
| # > 15 | 7 | 0 | 0 | 0 | 0 | 0 | 0 | 0 | 0 | 0 |
| Avg. | 160.4 | 2.7 | 2.8 | 2.3 | 2.4 | 2.5 | 3.0 | 3.0 | 2.7 | 2.0 |

**C (Wt)**

| | 100 | 200 | 300 | 400 | 500 | 600 | 700 | 800 | 900 | 1000 |
|---|---|---|---|---|---|---|---|---|---|---|
| n | 4 | 23 | 12 | 10 | 10 | 7 | 3 | 2 | 2 | 1 |
| Min. | 197 | 1 | 1 | 2 | 1 | 1 | 1 | 2 | 1 | 2 |
| Max. | 296 | 8 | 7 | 4 | 4 | 4 | 4 | 2 | 2 | 2 |
| # > 15 | 4 | 0 | 0 | 0 | 0 | 0 | 0 | 0 | 0 | 0 |
| Avg. | 230.2 | 3.1 | 3.0 | 2.5 | 1.9 | 1.7 | 2.7 | 2.0 | 1.5 | 2.0 |

**D (*mrc1Δ*)**

| | 100 | 200 | 300 | 400 | 500 | 600 | 700 | 800 | 900 | 1000 |
|---|---|---|---|---|---|---|---|---|---|---|
| n | 1 | 1 | 1 | 1 | 1 | 1 | 2 | 1 | 1 | 1 |
| Min. | 10 | 4 | 4 | 4 | 3 | 3 | 2 | 2 | 2 | 2 |
| Max. | 0 | 0 | 0 | 0 | 0 | 0 | 0 | 0 | 0 | 0 |
| # > 15 | 1 | 1 | 1 | 1 | 1 | 1 | 2 | 1 | 1 | 1 |
| Avg. | 3.3 | 2.0 | 1.7 | 1.7 | 1.6 | 1.8 | 2.0 | 1.5 | 1.5 | 1.5 |

**E (*mrc1Δ*)**

| | 100 | 200 | 300 | 400 | 500 | 600 | 700 | 800 | 900 | 1000 |
|---|---|---|---|---|---|---|---|---|---|---|
| n | 84 | 34 | 15 | 6 | 4 | 3 | 3 | 2 | 2 | 1 |
| Min. | 1 | 1 | 1 | 1 | 1 | 1 | 1 | 1 | 1 | 0 |
| Max. | 13 | 5 | 4 | 4 | 3 | 3 | 2 | 2 | 1 | 0 |
| # > 15 | 0 | 0 | 0 | 0 | 0 | 0 | 0 | 0 | 0 | 0 |
| Avg. | 3.9 | 2.4 | 2.1 | 2.2 | 2.2 | 2.0 | 1.3 | 1.5 | 1.0 | 0.0 |

**F (*mrc1Δ*)**

| | 100 | 200 | 300 | 400 | 500 | 600 | 700 | 800 | 900 | 1000 |
|---|---|---|---|---|---|---|---|---|---|---|
| n | 77 | 39 | 21 | 16 | 9 | 6 | 3 | 2 | 2 | 1 |
| Min. | 1 | 1 | 1 | 1 | 1 | 1 | 1 | 2 | 1 | 1 |
| Max. | 20 | 7 | 6 | 3 | 3 | 3 | 3 | 2 | 2 | 1 |
| # > 15 | 2 | 0 | 0 | 0 | 0 | 0 | 0 | 0 | 0 | 0 |
| Avg. | 4.3 | 2.6 | 2.5 | 1.8 | 1.7 | 1.7 | 2.0 | 2.0 | 1.5 | 1.0 |

**G (*cds1 Δ*)**

| | 100 | 200 | 300 | 400 | 500 | 600 | 700 | 800 | 900 | 1000 |
|---|---|---|---|---|---|---|---|---|---|---|
| n | 67 | 42 | 30 | 25 | 16 | 16 | 13 | 9 | 7 | 7 |
| Min. | 1 | 1 | 1 | 1 | 1 | 1 | 1 | 1 | 1 | 1 |
| Max. | 15 | 10 | 8 | 7 | 6 | 4 | 3 | 3 | 2 | 2 |
| # > 15 | 0 | 0 | 0 | 0 | 0 | 0 | 0 | 0 | 0 | 0 |
| Avg. | 4.7 | 3.7 | 2.9 | 2.4 | 2.5 | 2.0 | 1.8 | 1.7 | 1.4 | 1.4 |

**H (*cds1 Δ*)**

| | 100 | 200 | 300 | 400 | 500 | 600 | 700 | 800 | 900 | 1000 |
|---|---|---|---|---|---|---|---|---|---|---|
| n | 32 | 28 | 22 | 13 | 11 | 11 | 8 | 5 | 4 | 4 |
| Min. | 1 | 1 | 1 | 1 | 1 | 1 | 1 | 1 | 2 | 2 |
| Max. | 10 | 8 | 8 | 6 | 4 | 4 | 4 | 3 | 3 | 3 |
| # > 15 | 0 | 0 | 0 | 0 | 0 | 0 | 0 | 0 | 0 | 0 |
| Avg. | 3.7 | 3.4 | 2.6 | 2.5 | 2.1 | 2.0 | 2.0 | 2.2 | 2.5 | 2.5 |

**I (*cds1 Δ*)**

| | 100 | 200 | 300 | 400 | 500 | 600 | 700 | 800 | 900 | 1000 |
|---|---|---|---|---|---|---|---|---|---|---|
| n | 77 | 61 | 47 | 39 | 31 | 26 | 23 | 22 | 21 | 17 |
| Min. | 1 | 1 | 1 | 1 | 1 | 1 | 1 | 1 | 1 | 2 |
| Max. | 14 | 13 | 7 | 6 | 5 | 5 | 5 | 4 | 3 | 3 |
| # > 15 | 0 | 0 | 0 | 0 | 0 | 0 | 0 | 0 | 0 | 0 |
| Avg. | 4.3 | 4.0 | 3.1 | 2.7 | 2.5 | 2.5 | 2.4 | 2.3 | 2.1 | 2.3 |

**Supplemental Figure 4:** Individual Replicates of Cdc45 Thresholding for Wt, *cds1Δ* and *mrc1Δ*.

SUPPLEMENTAL 4

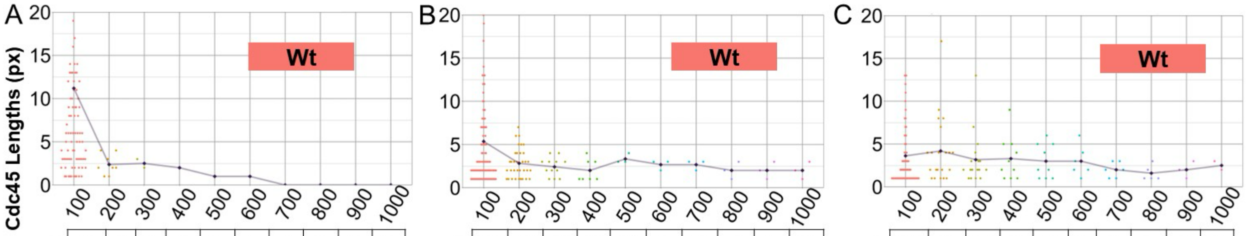


A

| | | | | | | | | | | |
|---|---|---|---|---|---|---|---|---|---|---|
| n | 94 | 8 | 2 | 1 | 1 | 1 | 1 | 1 | 1 | 1 |
| Min. | 1 | 1 | 2 | 2 | 1 | 1 | 0 | 0 | 0 | 0 |
| Max. | 55 | 4 | 3 | 2 | 1 | 1 | 0 | 0 | 0 | 0 |
| # > 20 | 18 | 0 | 0 | 0 | 0 | 0 | 0 | 0 | 0 | 0 |
| Avg. | 11.2 | 2.4 | 2.5 | 2.0 | 1.0 | 1.0 | 0.0 | 0.0 | 0.0 | 0.0 |

B

| | | | | | | | | | | |
|---|---|---|---|---|---|---|---|---|---|---|
| n | 139 | 34 | 12 | 8 | 3 | 3 | 3 | 3 | 3 | 3 |
| Min. | 1 | 1 | 1 | 1 | 3 | 2 | 2 | 1 | 1 | 1 |
| Max. | 59 | 7 | 4 | 4 | 4 | 3 | 3 | 3 | 3 | 3 |
| # > 20 | 7 | 0 | 0 | 0 | 0 | 0 | 0 | 0 | 0 | 0 |
| Avg. | 5.4 | 2.8 | 2.4 | 2.0 | 3.3 | 2.7 | 2.7 | 2.0 | 2.0 | 2.0 |

C

| | | | | | | | | | | |
|---|---|---|---|---|---|---|---|---|---|---|
| n | 117 | 21 | 17 | 10 | 9 | 6 | 6 | 5 | 3 | 2 |
| Min. | 1 | 1 | 1 | 1 | 1 | 1 | 1 | 1 | 1 | 2 |
| Max. | 37 | 17 | 13 | 9 | 6 | 6 | 3 | 3 | 3 | 3 |
| # > 20 | 4 | 0 | 0 | 0 | 0 | 0 | 0 | 0 | 0 | 0 |
| Avg. | 3.6 | 4.2 | 3.2 | 3.3 | 3.0 | 3.0 | 2.0 | 1.6 | 2.0 | 2.5 |

Cdc45 Thresholding Value →

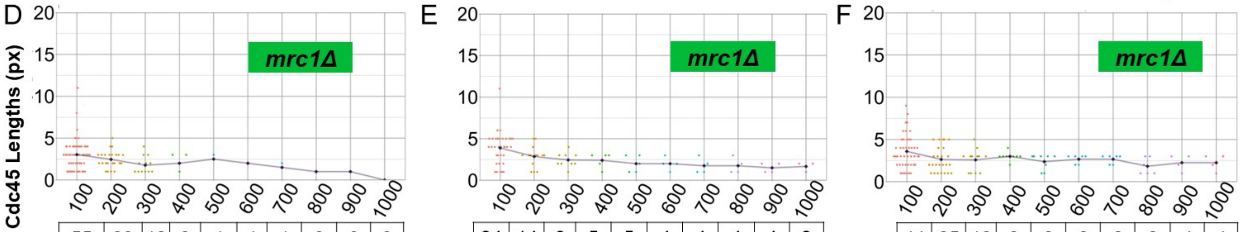


D

| | | | | | | | | | | |
|---|---|---|---|---|---|---|---|---|---|---|
| n | 55 | 23 | 12 | 3 | 1 | 1 | 1 | 0 | 0 | 0 |
| Min. | 1 | 1 | 1 | 1 | 3 | 2 | 2 | 0 | 0 | 0 |
| Max. | 11 | 5 | 4 | 3 | 3 | 2 | 2 | 0 | 0 | 0 |
| # > 20 | 0 | 0 | 0 | 0 | 0 | 0 | 0 | 0 | 0 | 0 |
| Avg. | 3.0 | 2.5 | 1.8 | 2.0 | 2.5 | 2.0 | 1.5 | 1.0 | 1.0 | 0.0 |

E

| | | | | | | | | | | |
|---|---|---|---|---|---|---|---|---|---|---|
| n | 34 | 14 | 9 | 5 | 5 | 4 | 4 | 4 | 4 | 3 |
| Min. | 1 | 1 | 1 | 1 | 1 | 1 | 1 | 1 | 1 | 1 |
| Max. | 11 | 5 | 4 | 3 | 3 | 3 | 3 | 3 | 2 | 2 |
| # >20 | 0 | 0 | 0 | 0 | 0 | 0 | 0 | 0 | 0 | 0 |
| Avg. | 3.9 | 2.9 | 2.4 | 2.4 | 2.0 | 2.0 | 1.8 | 1.8 | 1.5 | 1.7 |

F

| | | | | | | | | | | |
|---|---|---|---|---|---|---|---|---|---|---|
| n | 44 | 25 | 12 | 8 | 8 | 6 | 6 | 6 | 4 | 4 |
| Min. | 1 | 1 | 1 | 2 | 1 | 2 | 2 | 1 | 1 | 1 |
| Max. | 9 | 5 | 5 | 4 | 3 | 3 | 3 | 3 | 3 | 3 |
| # > 20 | 0 | 0 | 0 | 0 | 0 | 0 | 0 | 0 | 0 | 0 |
| Avg. | 3.6 | 2. | 2.6 | 3.0 | 2.4 | 2.7 | 2.7 | 1.8 | 2.2 | 2.2 |

Cdc45 Thresholding Value →

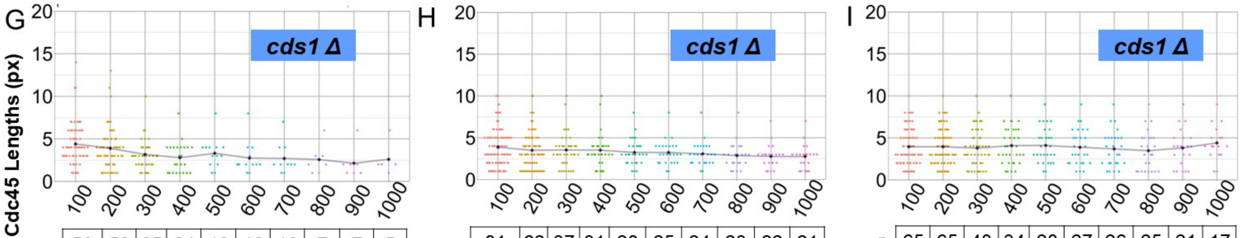


G

| | | | | | | | | | | |
|---|---|---|---|---|---|---|---|---|---|---|
| n | 59 | 53 | 35 | 24 | 13 | 12 | 10 | 7 | 7 | 5 |
| Min. | 1 | 1 | 1 | 1 | 1 | 1 | 1 | 1 | 1 | 1 |
| Max. | 14 | 13 | 10 | 8 | 8 | 8 | 7 | 6 | 6 | 6 |
| # > 20 | 0 | 0 | 0 | 0 | 0 | 0 | 0 | 0 | 0 | 0 |
| Avg. | 4.4 | 3.9 | 3.2 | 2.8 | 3.3 | 2.8 | 2.7 | 2.6 | 2.1 | 2.6 |

H

| | | | | | | | | | | |
|---|---|---|---|---|---|---|---|---|---|---|
| n | 81 | 62 | 37 | 31 | 28 | 25 | 24 | 23 | 22 | 21 |
| Min. | 1 | 1 | 1 | 1 | 1 | 1 | 1 | 1 | 1 | 1 |
| Max. | 10 | 10 | 10 | 10 | 8 | 8 | 8 | 8 | 7 | 7 |
| # > 20 | 0 | 0 | 0 | 0 | 0 | 0 | 0 | 0 | 0 | 0 |
| Avg. | 3.9 | 3.5 | 3.5 | 3.5 | 3.2 | 3.2 | 3.1 | 2.9 | 2.8 | 2.8 |

I

| | | | | | | | | | | |
|---|---|---|---|---|---|---|---|---|---|---|
| n | 65 | 65 | 48 | 34 | 28 | 27 | 26 | 25 | 21 | 17 |
| Min. | 1 | 1 | 1 | 1 | 1 | 1 | 1 | 1 | 1 | 1 |
| Max. | 10 | 10 | 10 | 10 | 9 | 9 | 9 | 9 | 9 | 9 |
| # > 20 | 0 | 0 | 0 | 0 | 0 | 0 | 0 | 0 | 0 | 0 |
| Avg. | 4.0 | 4.0 | 3.8 | 4.1 | 4.1 | 3.9 | 3.7 | 3.5 | 3.8 | 4.4 |

Cdc45 Thresholding Value →

**Supplemental Figure 5:** Individual Replicates of BrdU Smoothing for Wt, *cds1Δ* and *mrc1Δ*.

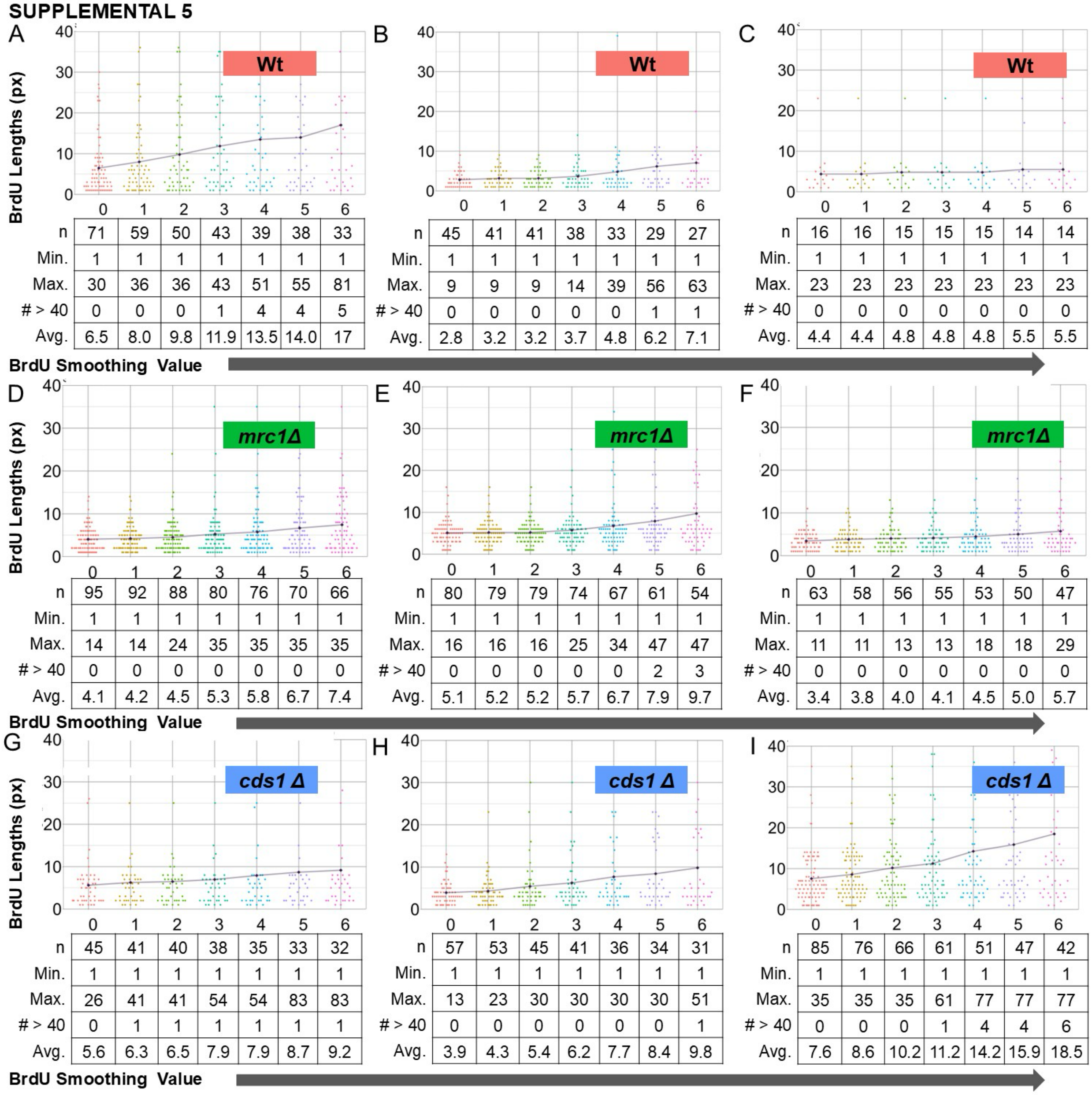


**A (Wt)**

| | 0 | 1 | 2 | 3 | 4 | 5 | 6 |
|---|---|---|---|---|---|---|---|
| n | 71 | 59 | 50 | 43 | 39 | 38 | 33 |
| Min. | 1 | 1 | 1 | 1 | 1 | 1 | 1 |
| Max. | 30 | 36 | 36 | 43 | 51 | 55 | 81 |
| # > 40 | 0 | 0 | 0 | 1 | 4 | 4 | 5 |
| Avg. | 6.5 | 8.0 | 9.8 | 11.9 | 13.5 | 14.0 | 17 |

**B (Wt)**

| | 0 | 1 | 2 | 3 | 4 | 5 | 6 |
|---|---|---|---|---|---|---|---|
| n | 45 | 41 | 41 | 38 | 33 | 29 | 27 |
| Min. | 1 | 1 | 1 | 1 | 1 | 1 | 1 |
| Max. | 9 | 9 | 9 | 14 | 39 | 56 | 63 |
| # > 40 | 0 | 0 | 0 | 0 | 0 | 1 | 1 |
| Avg. | 2.8 | 3.2 | 3.2 | 3.7 | 4.8 | 6.2 | 7.1 |

**C (Wt)**

| | 0 | 1 | 2 | 3 | 4 | 5 | 6 |
|---|---|---|---|---|---|---|---|
| n | 16 | 16 | 15 | 15 | 15 | 14 | 14 |
| Min. | 1 | 1 | 1 | 1 | 1 | 1 | 1 |
| Max. | 23 | 23 | 23 | 23 | 23 | 23 | 23 |
| # > 40 | 0 | 0 | 0 | 0 | 0 | 0 | 0 |
| Avg. | 4.4 | 4.4 | 4.8 | 4.8 | 4.8 | 5.5 | 5.5 |

**D (*mrc1Δ*)**

| | 0 | 1 | 2 | 3 | 4 | 5 | 6 |
|---|---|---|---|---|---|---|---|
| n | 95 | 92 | 88 | 80 | 76 | 70 | 66 |
| Min. | 1 | 1 | 1 | 1 | 1 | 1 | 1 |
| Max. | 14 | 14 | 24 | 35 | 35 | 35 | 35 |
| # > 40 | 0 | 0 | 0 | 0 | 0 | 0 | 0 |
| Avg. | 4.1 | 4.2 | 4.5 | 5.3 | 5.8 | 6.7 | 7.4 |

**E (*mrc1Δ*)**

| | 0 | 1 | 2 | 3 | 4 | 5 | 6 |
|---|---|---|---|---|---|---|---|
| n | 80 | 79 | 79 | 74 | 67 | 61 | 54 |
| Min. | 1 | 1 | 1 | 1 | 1 | 1 | 1 |
| Max. | 16 | 16 | 16 | 25 | 34 | 47 | 47 |
| # > 40 | 0 | 0 | 0 | 0 | 0 | 2 | 3 |
| Avg. | 5.1 | 5.2 | 5.2 | 5.7 | 6.7 | 7.9 | 9.7 |

**F (*mrc1Δ*)**

| | 0 | 1 | 2 | 3 | 4 | 5 | 6 |
|---|---|---|---|---|---|---|---|
| n | 63 | 58 | 56 | 55 | 53 | 50 | 47 |
| Min. | 1 | 1 | 1 | 1 | 1 | 1 | 1 |
| Max. | 11 | 11 | 13 | 13 | 18 | 18 | 29 |
| # > 40 | 0 | 0 | 0 | 0 | 0 | 0 | 0 |
| Avg. | 3.4 | 3.8 | 4.0 | 4.1 | 4.5 | 5.0 | 5.7 |

**G (*cds1 Δ*)**

| | 0 | 1 | 2 | 3 | 4 | 5 | 6 |
|---|---|---|---|---|---|---|---|
| n | 45 | 41 | 40 | 38 | 35 | 33 | 32 |
| Min. | 1 | 1 | 1 | 1 | 1 | 1 | 1 |
| Max. | 26 | 41 | 41 | 54 | 54 | 83 | 83 |
| # > 40 | 0 | 1 | 1 | 1 | 1 | 1 | 1 |
| Avg. | 5.6 | 6.3 | 6.5 | 7.9 | 7.9 | 8.7 | 9.2 |

**H (*cds1 Δ*)**

| | 0 | 1 | 2 | 3 | 4 | 5 | 6 |
|---|---|---|---|---|---|---|---|
| n | 57 | 53 | 45 | 41 | 36 | 34 | 31 |
| Min. | 1 | 1 | 1 | 1 | 1 | 1 | 1 |
| Max. | 13 | 23 | 30 | 30 | 30 | 30 | 51 |
| # > 40 | 0 | 0 | 0 | 0 | 0 | 0 | 1 |
| Avg. | 3.9 | 4.3 | 5.4 | 6.2 | 7.7 | 8.4 | 9.8 |

**I (*cds1 Δ*)**

| | 0 | 1 | 2 | 3 | 4 | 5 | 6 |
|---|---|---|---|---|---|---|---|
| n | 85 | 76 | 66 | 61 | 51 | 47 | 42 |
| Min. | 1 | 1 | 1 | 1 | 1 | 1 | 1 |
| Max. | 35 | 35 | 35 | 61 | 77 | 77 | 77 |
| # > 40 | 0 | 0 | 0 | 1 | 4 | 4 | 6 |
| Avg. | 7.6 | 8.6 | 10.2 | 11.2 | 14.2 | 15.9 | 18.5 |

**Supplemental Figure 6:** Individual Replicates of Rad51 Smoothing for Wt, *cds1Δ* and *mrc1Δ*.

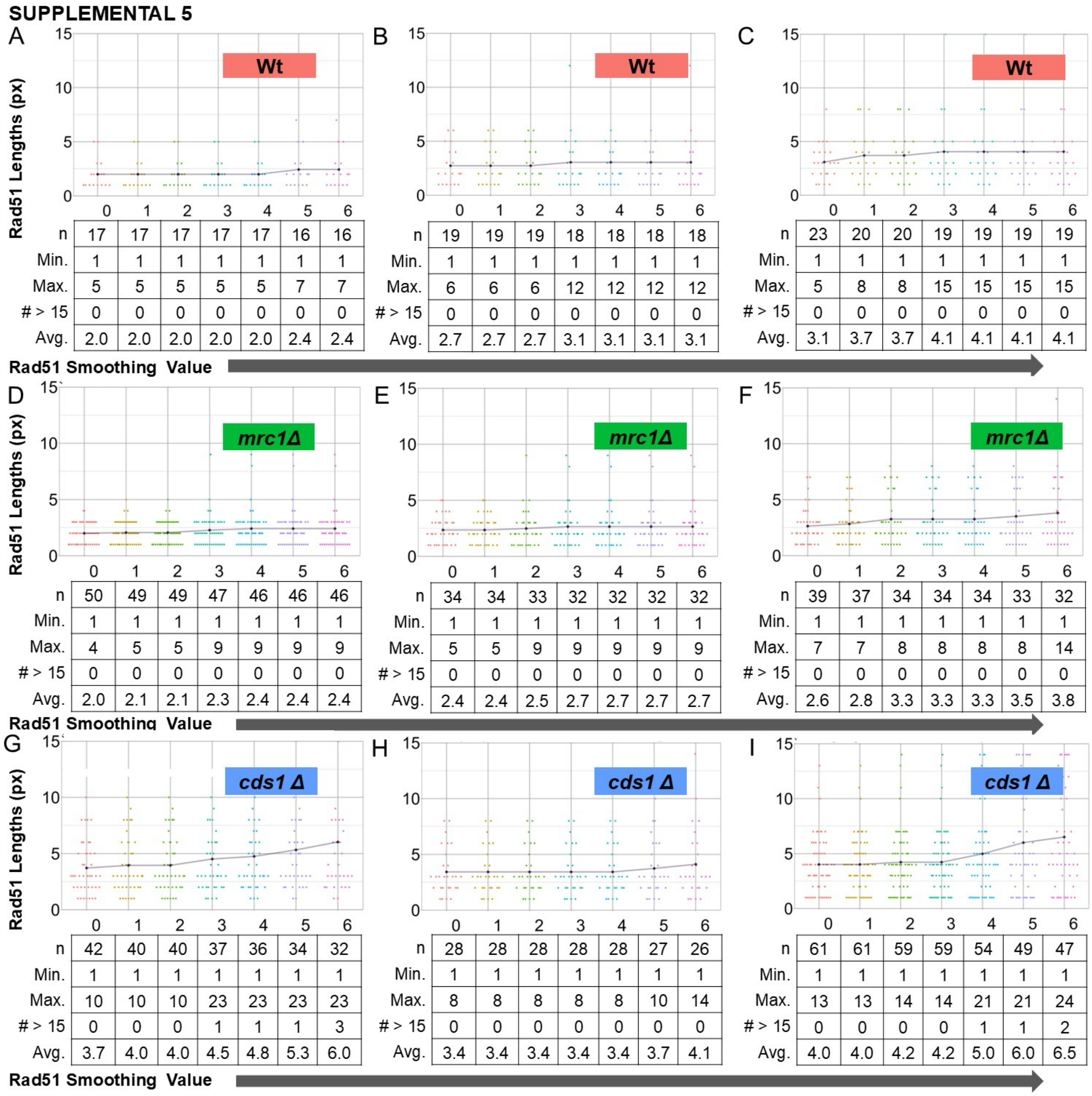


A (Wt)

| | 0 | 1 | 2 | 3 | 4 | 5 | 6 |
|---|---|---|---|---|---|---|---|
| n | 17 | 17 | 17 | 17 | 17 | 16 | 16 |
| Min. | 1 | 1 | 1 | 1 | 1 | 1 | 1 |
| Max. | 5 | 5 | 5 | 5 | 5 | 7 | 7 |
| # > 15 | 0 | 0 | 0 | 0 | 0 | 0 | 0 |
| Avg. | 2.0 | 2.0 | 2.0 | 2.0 | 2.0 | 2.4 | 2.4 |

B (Wt)

| | 0 | 1 | 2 | 3 | 4 | 5 | 6 |
|---|---|---|---|---|---|---|---|
| n | 19 | 19 | 19 | 18 | 18 | 18 | 18 |
| Min. | 1 | 1 | 1 | 1 | 1 | 1 | 1 |
| Max. | 6 | 6 | 6 | 12 | 12 | 12 | 12 |
| # > 15 | 0 | 0 | 0 | 0 | 0 | 0 | 0 |
| Avg. | 2.7 | 2.7 | 2.7 | 3.1 | 3.1 | 3.1 | 3.1 |

C (Wt)

| | 0 | 1 | 2 | 3 | 4 | 5 | 6 |
|---|---|---|---|---|---|---|---|
| n | 23 | 20 | 20 | 19 | 19 | 19 | 19 |
| Min. | 1 | 1 | 1 | 1 | 1 | 1 | 1 |
| Max. | 5 | 8 | 8 | 15 | 15 | 15 | 15 |
| # > 15 | 0 | 0 | 0 | 0 | 0 | 0 | 0 |
| Avg. | 3.1 | 3.7 | 3.7 | 4.1 | 4.1 | 4.1 | 4.1 |

D (*mrc1Δ*)

| | 0 | 1 | 2 | 3 | 4 | 5 | 6 |
|---|---|---|---|---|---|---|---|
| n | 50 | 49 | 49 | 47 | 46 | 46 | 46 |
| Min. | 1 | 1 | 1 | 1 | 1 | 1 | 1 |
| Max. | 4 | 5 | 5 | 9 | 9 | 9 | 9 |
| # > 15 | 0 | 0 | 0 | 0 | 0 | 0 | 0 |
| Avg. | 2.0 | 2.1 | 2.1 | 2.3 | 2.4 | 2.4 | 2.4 |

E (*mrc1Δ*)

| | 0 | 1 | 2 | 3 | 4 | 5 | 6 |
|---|---|---|---|---|---|---|---|
| n | 34 | 34 | 33 | 32 | 32 | 32 | 32 |
| Min. | 1 | 1 | 1 | 1 | 1 | 1 | 1 |
| Max. | 5 | 5 | 9 | 9 | 9 | 9 | 9 |
| # > 15 | 0 | 0 | 0 | 0 | 0 | 0 | 0 |
| Avg. | 2.4 | 2.4 | 2.5 | 2.7 | 2.7 | 2.7 | 2.7 |

F (*mrc1Δ*)

| | 0 | 1 | 2 | 3 | 4 | 5 | 6 |
|---|---|---|---|---|---|---|---|
| n | 39 | 37 | 34 | 34 | 34 | 33 | 32 |
| Min. | 1 | 1 | 1 | 1 | 1 | 1 | 1 |
| Max. | 7 | 7 | 8 | 8 | 8 | 8 | 14 |
| # > 15 | 0 | 0 | 0 | 0 | 0 | 0 | 0 |
| Avg. | 2.6 | 2.8 | 3.3 | 3.3 | 3.3 | 3.5 | 3.8 |

G (*cds1 Δ*)

| | 0 | 1 | 2 | 3 | 4 | 5 | 6 |
|---|---|---|---|---|---|---|---|
| n | 42 | 40 | 40 | 37 | 36 | 34 | 32 |
| Min. | 1 | 1 | 1 | 1 | 1 | 1 | 1 |
| Max. | 10 | 10 | 10 | 23 | 23 | 23 | 23 |
| # > 15 | 0 | 0 | 0 | 1 | 1 | 1 | 3 |
| Avg. | 3.7 | 4.0 | 4.0 | 4.5 | 4.8 | 5.3 | 6.0 |

H (*cds1 Δ*)

| | 0 | 1 | 2 | 3 | 4 | 5 | 6 |
|---|---|---|---|---|---|---|---|
| n | 28 | 28 | 28 | 28 | 28 | 27 | 26 |
| Min. | 1 | 1 | 1 | 1 | 1 | 1 | 1 |
| Max. | 8 | 8 | 8 | 8 | 8 | 10 | 14 |
| # > 15 | 0 | 0 | 0 | 0 | 0 | 0 | 0 |
| Avg. | 3.4 | 3.4 | 3.4 | 3.4 | 3.4 | 3.7 | 4.1 |

I (*cds1 Δ*)

| | 0 | 1 | 2 | 3 | 4 | 5 | 6 |
|---|---|---|---|---|---|---|---|
| n | 61 | 61 | 59 | 59 | 54 | 49 | 47 |
| Min. | 1 | 1 | 1 | 1 | 1 | 1 | 1 |
| Max. | 13 | 13 | 14 | 14 | 21 | 21 | 24 |
| # > 15 | 0 | 0 | 0 | 0 | 1 | 1 | 2 |
| Avg. | 4.0 | 4.0 | 4.2 | 4.2 | 5.0 | 6.0 | 6.5 |

**Supplemental Figure 7:** Individual Replicates of Cdc45 Smoothing iteration.

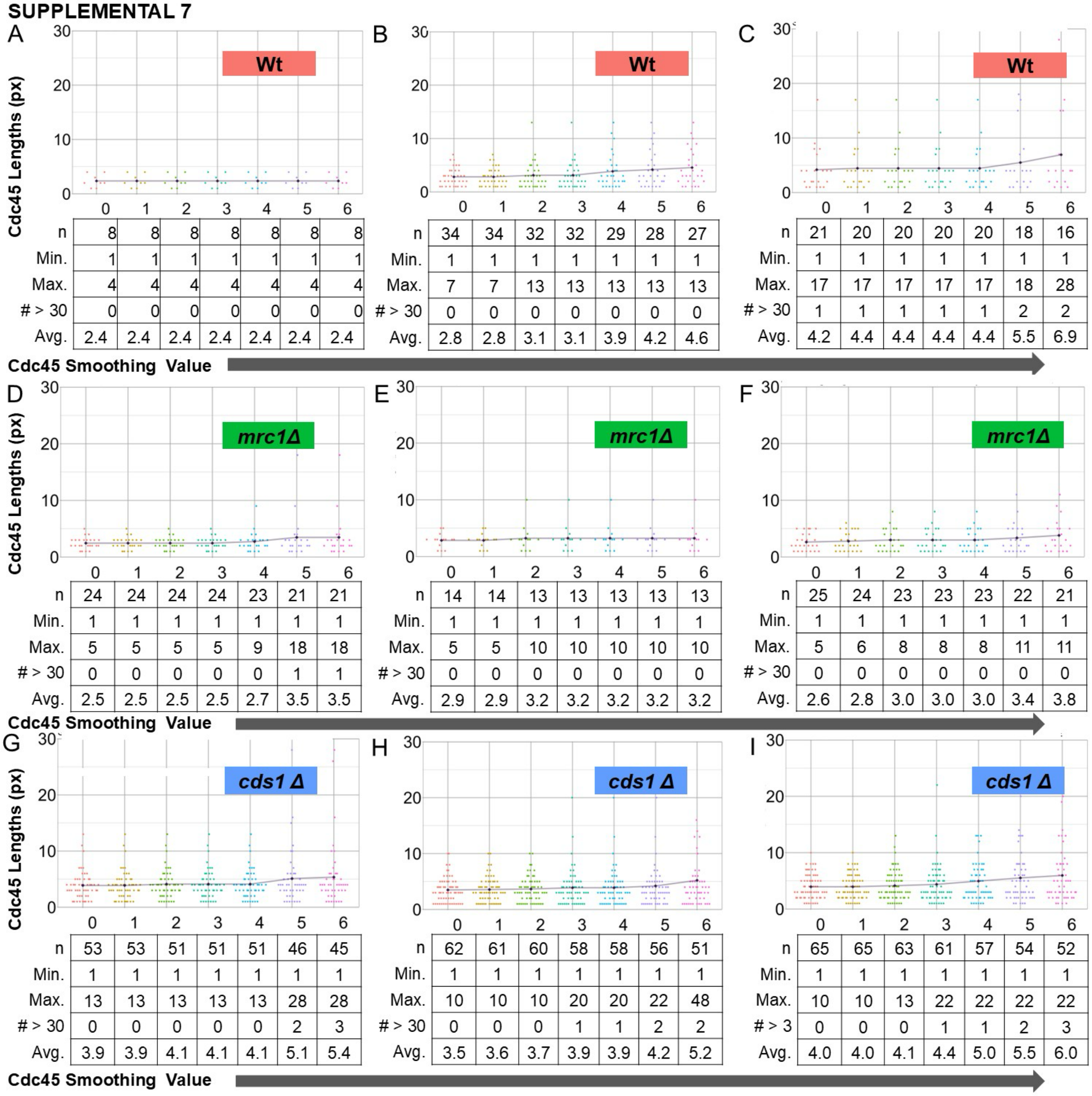


A (Wt)

| Cdc45 Smoothing Value | 0 | 1 | 2 | 3 | 4 | 5 | 6 |
|---|---|---|---|---|---|---|---|
| n | 8 | 8 | 8 | 8 | 8 | 8 | 8 |
| Min. | 1 | 1 | 1 | 1 | 1 | 1 | 1 |
| Max. | 4 | 4 | 4 | 4 | 4 | 4 | 4 |
| # > 30 | 0 | 0 | 0 | 0 | 0 | 0 | 0 |
| Avg. | 2.4 | 2.4 | 2.4 | 2.4 | 2.4 | 2.4 | 2.4 |

B (Wt)

| Cdc45 Smoothing Value | 0 | 1 | 2 | 3 | 4 | 5 | 6 |
|---|---|---|---|---|---|---|---|
| n | 34 | 34 | 32 | 32 | 29 | 28 | 27 |
| Min. | 1 | 1 | 1 | 1 | 1 | 1 | 1 |
| Max. | 7 | 7 | 13 | 13 | 13 | 13 | 13 |
| # > 30 | 0 | 0 | 0 | 0 | 0 | 0 | 0 |
| Avg. | 2.8 | 2.8 | 3.1 | 3.1 | 3.9 | 4.2 | 4.6 |

C (Wt)

| Cdc45 Smoothing Value | 0 | 1 | 2 | 3 | 4 | 5 | 6 |
|---|---|---|---|---|---|---|---|
| n | 21 | 20 | 20 | 20 | 20 | 18 | 16 |
| Min. | 1 | 1 | 1 | 1 | 1 | 1 | 1 |
| Max. | 17 | 17 | 17 | 17 | 17 | 18 | 28 |
| # > 30 | 1 | 1 | 1 | 1 | 1 | 2 | 2 |
| Avg. | 4.2 | 4.4 | 4.4 | 4.4 | 4.4 | 5.5 | 6.9 |

D (*mrc1Δ*)

| Cdc45 Smoothing Value | 0 | 1 | 2 | 3 | 4 | 5 | 6 |
|---|---|---|---|---|---|---|---|
| n | 24 | 24 | 24 | 24 | 23 | 21 | 21 |
| Min. | 1 | 1 | 1 | 1 | 1 | 1 | 1 |
| Max. | 5 | 5 | 5 | 5 | 9 | 18 | 18 |
| # > 30 | 0 | 0 | 0 | 0 | 0 | 1 | 1 |
| Avg. | 2.5 | 2.5 | 2.5 | 2.5 | 2.7 | 3.5 | 3.5 |

E (*mrc1Δ*)

| Cdc45 Smoothing Value | 0 | 1 | 2 | 3 | 4 | 5 | 6 |
|---|---|---|---|---|---|---|---|
| n | 14 | 14 | 13 | 13 | 13 | 13 | 13 |
| Min. | 1 | 1 | 1 | 1 | 1 | 1 | 1 |
| Max. | 5 | 5 | 10 | 10 | 10 | 10 | 10 |
| # > 30 | 0 | 0 | 0 | 0 | 0 | 0 | 0 |
| Avg. | 2.9 | 2.9 | 3.2 | 3.2 | 3.2 | 3.2 | 3.2 |

F (*mrc1Δ*)

| Cdc45 Smoothing Value | 0 | 1 | 2 | 3 | 4 | 5 | 6 |
|---|---|---|---|---|---|---|---|
| n | 25 | 24 | 23 | 23 | 23 | 22 | 21 |
| Min. | 1 | 1 | 1 | 1 | 1 | 1 | 1 |
| Max. | 5 | 6 | 8 | 8 | 8 | 11 | 11 |
| # > 30 | 0 | 0 | 0 | 0 | 0 | 0 | 0 |
| Avg. | 2.6 | 2.8 | 3.0 | 3.0 | 3.0 | 3.4 | 3.8 |

G (*cds1 Δ*)

| Cdc45 Smoothing Value | 0 | 1 | 2 | 3 | 4 | 5 | 6 |
|---|---|---|---|---|---|---|---|
| n | 53 | 53 | 51 | 51 | 51 | 46 | 45 |
| Min. | 1 | 1 | 1 | 1 | 1 | 1 | 1 |
| Max. | 13 | 13 | 13 | 13 | 13 | 28 | 28 |
| # > 30 | 0 | 0 | 0 | 0 | 0 | 2 | 3 |
| Avg. | 3.9 | 3.9 | 4.1 | 4.1 | 4.1 | 5.1 | 5.4 |

H (*cds1 Δ*)

| Cdc45 Smoothing Value | 0 | 1 | 2 | 3 | 4 | 5 | 6 |
|---|---|---|---|---|---|---|---|
| n | 62 | 61 | 60 | 58 | 58 | 56 | 51 |
| Min. | 1 | 1 | 1 | 1 | 1 | 1 | 1 |
| Max. | 10 | 10 | 10 | 20 | 20 | 22 | 48 |
| # > 30 | 0 | 0 | 0 | 1 | 1 | 2 | 2 |
| Avg. | 3.5 | 3.6 | 3.7 | 3.9 | 3.9 | 4.2 | 5.2 |

I (*cds1 Δ*)

| Cdc45 Smoothing Value | 0 | 1 | 2 | 3 | 4 | 5 | 6 |
|---|---|---|---|---|---|---|---|
| n | 65 | 65 | 63 | 61 | 57 | 54 | 52 |
| Min. | 1 | 1 | 1 | 1 | 1 | 1 | 1 |
| Max. | 10 | 10 | 13 | 22 | 22 | 22 | 22 |
| # > 3 | 0 | 0 | 0 | 1 | 1 | 2 | 3 |
| Avg. | 4.0 | 4.0 | 4.1 | 4.4 | 5.0 | 5.5 | 6.0 |

for Wt, *cds1Δ* and *mrc1Δ*.

**Supplemental Figure 8: The effect of iterative window sizing on Rad51 and Cdc45 distribution near forks.**

A) Rad51 located with forks increases in most in *mrc1*Δ, suggesting that *mrc1*Δ forks accumulate Rad51 that spreads from the fork into unreplicated areas. In contrast, *cds1*Δ forks retain the most Rad51 in unreplicated areas that are away from forks, suggesting DNA damage and more spread away from collapsing forks. From left to right, pixels into the replicated fork area increase from 2 to 10. Extending the number of fork-pixels into replicated area has a small effect on Rad51 localization, regardless of genotype. The 10x10 extreme situation (bottom right) is dominated by the effect of un-replicated pixels made fork-proximal.

B) Expanded window-size effects on Cdc45 colocalization for *cds1*Δ *(pink), mrc1*Δ *(green),* and wild type (blue). With no pixels included on either side of the tip, Cdc45 is primarily unreplicated ("COLO UNR REGION") regardless of fork window around BrdU tips. As the grid goes from top to bottom, the number of pixels extending in the unreplicated area near the fork increases from 2 to 10.
